# De novo Design of Translational RNA Repressors

**DOI:** 10.1101/501767

**Authors:** Paul D. Carlson, Cameron J. Glasscock, Julius B. Lucks

## Abstract

A central goal of synthetic biology is the development of methods for the predictable control of gene expression. RNA is an attractive substrate by which to achieve this goal because the relationship between its sequence, structure, and function is being uncovered with increasing depth. In addition, design approaches that use this relationship are becoming increasingly effective, as evidenced by significant progress in the *de novo* design of RNA-based gene regulatory mechanisms that activate transcription and translation in bacterial cells. However, the design of synthetic RNA mechanisms that are efficient and versatile repressors of gene expression has lagged, despite their importance for gene regulation and genetic circuit construction. We address this gap by developing two new classes of RNA regulators, toehold repressors and looped antisense oligonucleotides (LASOs), that repress translation of a downstream gene in response to an arbitrary input RNA sequence. Characterization studies show that these designed RNAs robustly repress translation, are highly orthogonal, and can be multiplexed with translational activators. We show that our LASO design can repress endogenous mRNA targets and distinguish between closely-related genes with a high degree of specificity and predictability. These results demonstrate significant yet easy-to-implement improvements in the design of synthetic RNA repressors for synthetic biology, and point more broadly to design principles for repressive RNA interactions relevant to modern drug design.

## INTRODUCTION

RNAs play myriad functional roles in the cell across all domains of life^1^. In prokaryotes, many regulatory non-coding RNAs (ncRNAs) act within the 5’ untranslated region (5’ UTR) of the gene they control to regulate the processes of transcription, translation, and mRNA degradation^2-4^. In nature, these RNAs can respond to a variety of signals, including ions^5^, metabolites^3,6^, temperature changes^7^, proteins^8^, and other regulatory RNAs^2,9,10^, effectively acting as important regulatory sensors of the cellular environment. Given the widespread distribution and important functional roles of these RNAs in nature^3^, they have been important targets for RNA engineering efforts.

Two important classes of ncRNAs are *trans*-acting bacterial antisense RNAs (asRNAs) and small RNAs (sRNAs), which act through direct base-pairing to the regulated mRNA to control downstream gene expression^11^. This often occurs through the binding of the *trans*-acting RNA to its target sequence, which is typically a portion of the 5’ untranslated region (5’ UTR) or coding sequence that includes the ribosome binding site (RBS) or start codon of the gene to inhibit its translation initiation^12^. Antisense RNAs are typically countertranscripts on the order of hundreds of nucleotides in length that are the perfect reverse-complement of their target. Conversely, sRNAs are often expressed from different genomic loci than their target, tend to be shorter than asRNAs, and often include Hfq protein binding scaffold sequences to facilitate Hfq-mediated sRNA-mRNA interactions^13,14^.

The relatively simple mode of action of asRNAs and sRNAs made them attractive for early RNA engineering efforts. For example, Soloman et al.^15^ used an asRNA to control glucokinase expression, achieving a 25% reduction in Glk activity. Similarly, libraries of sRNA translational repressors have been leveraged for a variety of applications^16^, including to knockdown gene expression in *E. coli* to improve titers of desired compounds produced from metabolic pathways^17^, and to create combinatorial screening libraries of sRNAs for the fine-tuned, simultaneous repression of multiple targets^18^. Moreover, studies of sRNA-target interactions have begun to uncover design rules for translational repressors^17,19^, making asRNAs and sRNAs key components of the RNA synthetic biology toolbox.

While progress in developing and applying asRNAs and sRNAs has been made, several key challenges have remained. First, repression of the target gene is often modest, with average repression levels reported in the range of 25–60%^15,19^. Second, the effects of Hfq are variable depending on the expression levels of individual RNA species and the Hfq-binding scaffold employed in the sRNA design^19,20^, creating uncertainty in sRNA design principles that have necessitated screening large numbers of designs to find regulators that function properly. Third, since asRNAs are typically designed to bind across the translation start site, there could be unintended off-target repression effects due to the sequence similarities between the Shine-Dalgarno sequence and AUG start codon of the intended target of an asRNA and non-cognate mRNAs^19,21^. Finally, intramolecular secondary structures formed by the asRNA or target can inhibit bimolecular hybridization and thus regulatory function^15^, and the effects of these inhibitory structures are not fully considered in synthetic asRNA design rules^19,21^.

These challenges have started to be addressed in recent years with the development of synthetic RNA regulatory mechanisms, or riboregulators. The first example of a naturally-inspired, synthetic riboregulator consists of a stem-loop structure that occludes the ribosome binding site (RBS) of a downstream gene, preventing translation^22^. Binding of a *trans*-acting RNA opens the stem-loop to allow downstream translation^23^, creating a synthetic off-to-on translation activation switch. Later work showed that orthogonal variants of this system could serve as a genetic “switchboard” to independently control different metabolic pathways in *E. coli*^24^. Building off of this concept, the capabilities of synthetic RNA activators have expanded even further with the development of the Nucleic Acid Package (NUPACK)^25,26^, which has enabled the *de novo* design of new types of riboregulators. Specifically, translational activators called toehold switches have been generated using NUPACK to design RNA sequences that fold into structural motifs that are modifications of the original riboregulatory scaffold, and that regulate their targets with large dynamic ranges with no sequence restrictions on the input RNA species^27^. This innovation has enabled the creation of large libraries of orthogonal toehold switch regulators^27^, inexpensive diagnostics to detect nucleic acids^28-30^, and complex multi-input logic devices^31^. NUPACK has also been used to design small transcription activating RNAs (STARs), which activate gene expression at the level of transcription and can serve as the basis for RNA-only logic gates and circuitry^32,33^. Interestingly, while progress has been made in the activation of gene expression using designed synthetic RNA systems, the development of similarly versatile RNA repressors of gene expression has lagged.

In this work, we address this shortcoming and devise methods to bridge the gap between synthetic RNA activators and repressors by developing strategies for the *de novo* design of repressive translational riboregulators called toehold repressors. Using designed sequences to invert the structural changes of toehold activators, we created a new class of toehold repressors that robustly repress downstream gene expression in response to an RNA input of arbitrary sequence. In the process, we realized that the same design elements of toehold translational repressors could be applied to asRNAs and sRNAs, which led us to create a new type of RNA translational repressor, called a looped antisense oligonucleotide (LASO). LASOs overcome the design challenges of asRNAs and sRNAs to create a regulator that can strongly repress a gene of interest with minimal off-target interactions that can be predicted using simple free energy considerations. Additionally, LASOs are compatible with RBS libraries, making them useful for genetic network optimization. We anticipate that the development of the LASO concept could have implications for RNA-mediated gene regulation in a variety of organisms, and may inform the design of new classes of oligonucleotide therapies^34,35^.

## RESULTS

### NUPACK can be used to design toehold repressors

Previous studies have shown that NUPACK can be used to design translational activators called toehold switches (Figure 1A)^27,31^. The core element of this design is an RNA hairpin that sequesters the RBS and start codon of a downstream gene to inhibit translation. By adding a trigger RNA that is designed to bind to the upstream portion of this hairpin, the hairpin can be opened to free the RBS and start codon to initiate translation. This trigger-target interaction is seeded at a single-stranded region called a toehold. The emphasis of this design on sequestering the RBS inside a hairpin loop, rather than through direct base pairing interactions, creates a flexible design space to enable the construction orthogonal (non-interacting) switches with large dynamic ranges^27^. These designs can be further extended to perform complex cellular logic^31^. Given these advantages, we sought to leverage the architecture of this design to develop a translational repressor that allows for translation of a downstream gene in the absence of any external factors, and is repressed in the presence of a corresponding trigger RNA.

**Figure 1.**
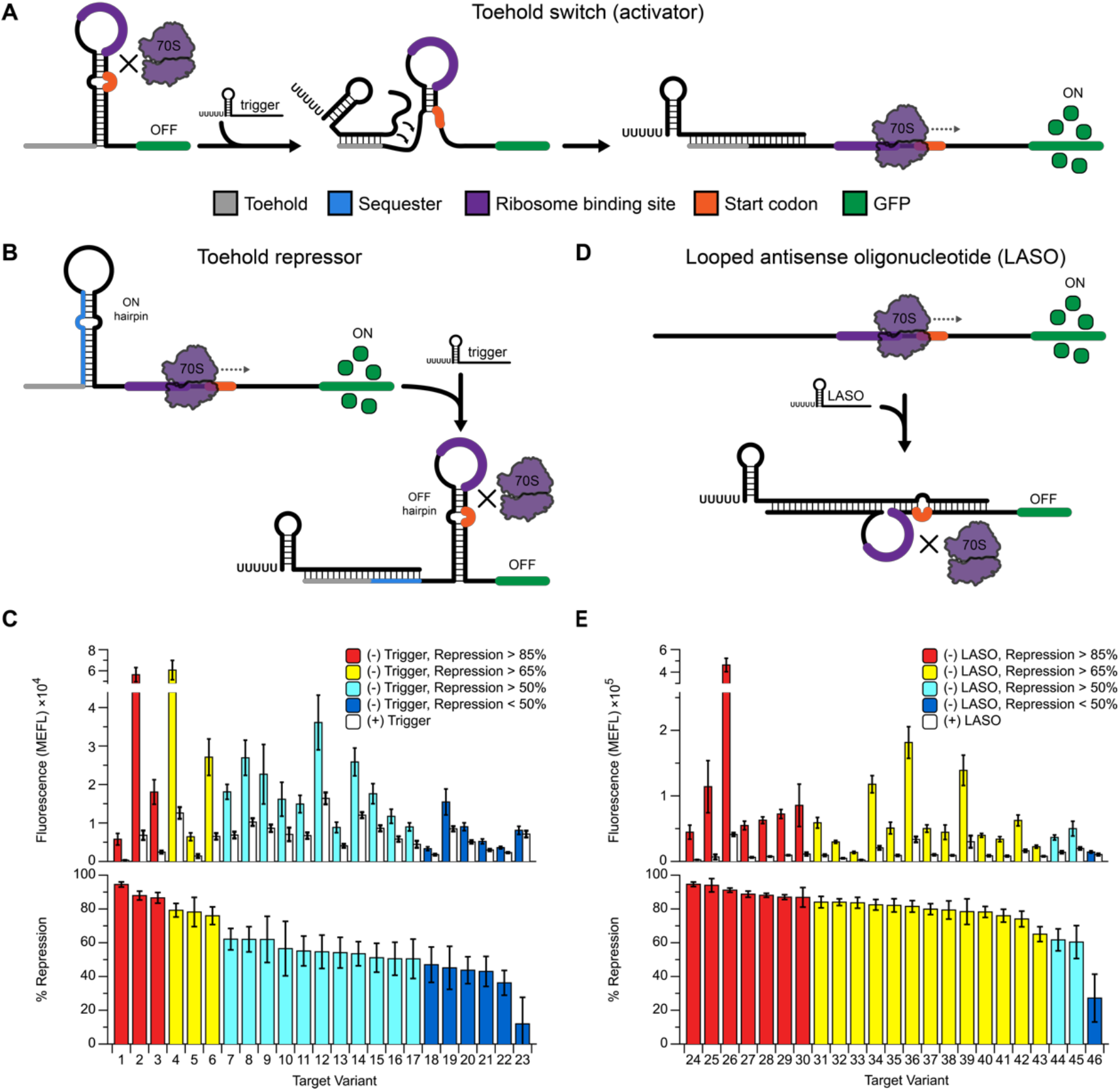
Design and performance of translational RNA regulators. (A) The toehold switch^27^ is a *de novo*-designed translational activator. Mechanistic model: the ribosome binding site (purple) and start codon (orange) of a downstream reporter gene (green) are constrained within a hairpin structure, inhibiting translation. Binding of an antisense trigger RNA opens the hairpin by toehold-mediated strand displacement (grey), freeing the translation start site and allowing expression of the reporter. (B) Proposed strategy to create a repressive version of the toehold switch, called a toehold repressor. Addition of a designed upstream sequester sequence (blue) converts the default toehold switch OFF state into an ON state by creating an alternative structure that leaves the RBS and start codon accessible. Trigger binding to the toehold and sequester regions opens the ON state hairpin, allowing the OFF hairpin to form and repress translation. (C) Performance of 23 designed toehold repressor target-trigger pairs with ON and OFF fluorescence (top) and percent repression (100% – OFF/ON) (bottom). (D) An alternative strategy to create translational repressors, termed a Looped Antisense Oligonucleotide (LASO). In this approach, the LASO binds directly around the translation start site of the target without binding directly to the ribosome binding site or the start codon. (E) Performance of 23 LASO-target pairs from (D) with ON and OFF fluorescence (top) and percent repression (bottom). Error bars represent at least 6 biological replicates collected on two separate days.

Our first translational repressor design, which we call a toehold repressor, is an RNA sequence within the 5’ UTR of a gene to be controlled that is designed to fold into two competing hairpin structures (Figure 1B). The minimum free energy structure consists of a 15nt single stranded toehold region followed by a hairpin (ON hairpin) that is formed upstream of a single stranded domain containing the ribosome binding site (RBS) and start codon. By itself, this structure allows for active translation of the downstream reporter, giving a default ON state to the regulator. Interaction with a designed trigger that is complementary to the toehold region and a portion of the ON hairpin is designed to cause the ON hairpin to undergo a toehold-mediated strand displacement, allowing an alternative hairpin structure (OFF hairpin) to form. The OFF hairpin is designed to form around the RBS and start codon to inhibit translation initiation much like in the case of toehold switches. Importantly, the RBS and start codon are not designed to directly base-pair with any other region of the repressor, resulting in no hard sequence constraints on the design.

A set of 23 toehold repressors was designed in NUPACK using two variations of this overall design approach. For 11 of the designs, we used domain lengths such that both hairpins have the same structure as the original toehold switch design^27^ (Supplementary Figure 1A). For the other 12 designs, we designed the OFF hairpin structure to have a smaller loop and elongated stem, which has been shown to have low leak^28^ (Supplementary Figure 1B). Additionally, for this design, the ON hairpin is designed with a longer, more stable, stem with the goal of thermodynamically driving the formation of the ON structure in the absence of a trigger RNA.

To test the toehold repressor designs, NUPACK-designed sequences were cloned in between the J23101 promoter^36^ and the superfolder GFP (sfGFP) coding sequence followed by a TrrnB transcriptional terminator in a medium copy plasmid (target plasmid, Supplementary Figure 2A). Corresponding trigger sequences were cloned in between a J23101 promoter and T500 transcriptional terminator on a high copy plasmid (trigger plasmid, Supplementary Figure 2B, see Materials and Methods). Target plasmids were transformed into *E. coli* TG1 with the corresponding trigger plasmid or a control plasmid (pJBL002), which expresses a fragment of the *rrnB* operon^37^. Individual colonies were picked, grown overnight in LB, diluted 1:50 in M9 minimal media, grown for 6h, and then characterized for sfGFP expression by flow cytometry (see Materials and Methods). All but one of the designs showed a significant decrease in fluorescence upon addition of the trigger plasmid, with the best design giving 94% (±1.5%) repression (Figure 1C).

### Designed trans-acting antisense RNAs can repress translation without binding directly to the RBS or start codon

We next sought to assess whether a simpler design approach could be used to achieve translational repression. Previous work has developed antisense RNAs (asRNAs) and small RNAs (sRNAs) as platforms for RNA-based translational repression^15,17,19,21^. Both mechanisms work by binding at or near the translation start site of their target: asRNAs often interfere with translation initiation by binding directly to the RBS and/or start codon of their target mRNA^15^, while sRNAs often utilize the protein co-factor Hfq to facilitate sRNA-target interactions. While powerful, both have limitations. For designed asRNAs, the conserved nature of RBS and start codon sequences^38,39^ could create unintended repression of off-target genes that share similar regulatory sequences^19,21^. For engineered sRNAs, the scaffold sequences used to recruit Hfq to facilitate sRNA-target interactions can have unpredictable effects depending on the scaffold used or the level of expression of the sRNA^19,20^. The toehold switch and toehold repressor designs both address these shortcomings by sequestering the RBS and start codon in looped regions to remove sequence constraints and by not requiring RNA-protein interactions to function. However, they add structural complexity beyond the simple complementarity rules of asRNA design. We therefore sought to merge the best of each approach and create a modified asRNA design that utilizes simple complementarity to the target region, but places the RBS and start codon in a looped context to free hard sequence constraints.

The looped antisense oligonucleotide (LASO) design concept that follows this strategy is outlined in Figure 1D. In this strategy, the 5’ UTR of the target RNA has no design constraints beyond being sufficiently unstructured to facilitate active translation. The LASO RNA is designed to bind across the translation start site while avoiding direct base-pairing to the RBS and start codon. The bound structure of this target/LASO complex therefore resembles the OFF hairpin of a toehold switch or toehold repressor, but with formation occurring in *trans* rather than in *cis*.

To test the LASO strategy, we designed 23 distinct target regions and placed them into the same expression context as our previous designs. To ensure active translation from these sequences, we started with toehold switch designs from Green et al.^27^ and removed the 5’ side of the hairpin structure, resulting in a library of unstructured and translationally-active sequences. We then designed LASOs to bind around the translation start site as shown in Supplementary Figure 3. Each design was tested in the same manner as the toehold repressors, with every design except one showing greater than 50% repression, and seven designs giving greater than 85% repression. The best-performing design showed 95% (±1.3%) repression, with median repression surpassing that of the toehold repressor designs (Figure 1E, Supplementary Figure 4). These data demonstrate that designed RNAs can efficiently repress translation without an Hfq binding site and without binding directly to the RBS or start codon.

### Translational repressors can form orthogonal libraries

An important quality of engineered riboregulators is the ability to create orthogonal (non-interacting) sets such that each trigger only represses its cognate target with little repression of non-cognate targets^24,27,40,41^. Such orthogonality of regulators is a requirement for their use in more sophisticated genetic networks^31^. To identify orthogonal sets from our pool of 46 toehold and LASO repressor designs, we first identified the 24 best-performing designs from the pool (Figure 1C,E), and then characterized expression from every target/trigger combination using flow cytometry (Figure 2, Supplementary Figure 5). By defining weak cross-talk as 20% off-target repression measured with respect to the no-trigger control^41^, we were able to identify a set of 10 mutually orthogonal trigger/LASO and target pairs, all with >70% repression of the cognate pair (Figure 2C). This demonstrates that our translational repressor designs can form sets of regulators with high on-target and low off-target repression, which could serve as the basis for more complex RNA circuitry.

**Figure 2.**
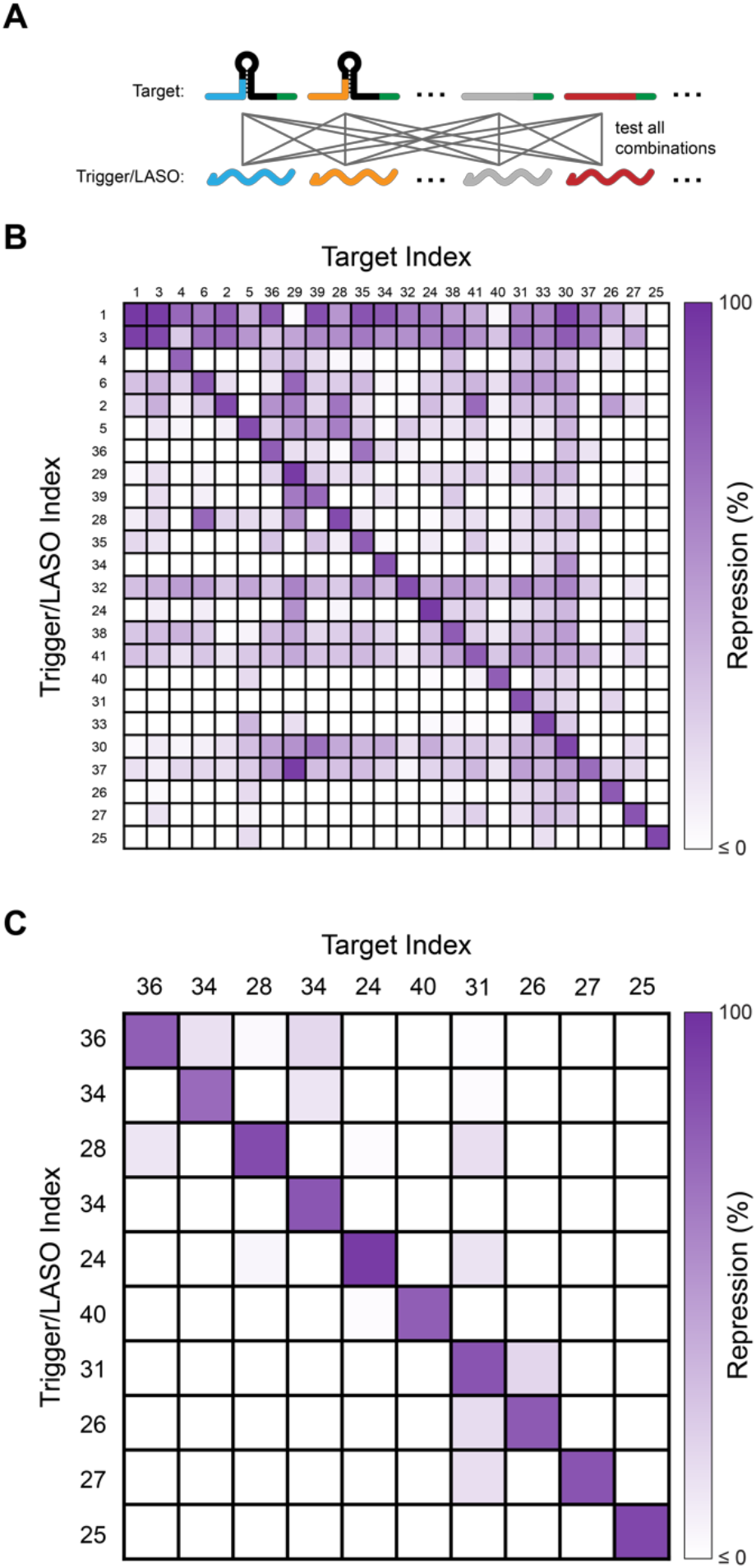
Assessing the orthogonality of translational repressors. (A) To test crosstalk in a set of designs, the expression of cells harbouring all possible combinations of plasmids encoding target and trigger/LASO expression constructs are measured and compared to the expression of each target plasmid expressed with a no-trigger control plasmid. (B) 24×24 orthogonality matrix of the best-performing designs from Figure 1. Each portion of the matrix corresponds to the repression observed from that combination of target and trigger/LASO expression plasmids compared to the no-trigger control. (C) A 10×10 subset of the designs tested in (B) selected to be mutually orthogonal, defined as showing less than 20% repression from all non-cognate pairs.

### Rational RNA engineering strategies improve the dynamic range of toehold repressors

Having established that we could create libraries of orthogonal regulators, we next pursued RNA engineering efforts to improve the performance of our toehold repressor design. An interesting trend we observed was a higher average ON level among the LASO targets compared to the toehold repressor library (Figure 3A). We suspected that the strong hairpin 4nt upstream of the RBS in the toehold repressor target designs may inhibit expression in the ON state, requiring the ribosome to partially unwind the base of the hairpin before initiating translation (Figure 3B, Supplementary Figure 1). Indeed, previous work demonstrated that separating a RBS and a strong upstream hairpin with 8nt or 12nt spacers can substantially increase translation compared to only a 4nt spacer^42^. To test this hypothesis, we added additional spacer sequences (+4nt and +8nt) to four toehold repressors with low ON levels (Figure 3C). In all cases, the ON level increased significantly, with the highest fluorescence observed when the hairpin-RBS spacer was increased to 12nt. For the best-performing design (design #1), the dynamic range (ON fluorescence divided by OFF fluorescence) increased from 20-fold to 31-fold (Figure 3D,E). Increases in dynamic range were also observed for designs 5, 13, and 18 (Supplementary Figure 6).

**Figure 3.**
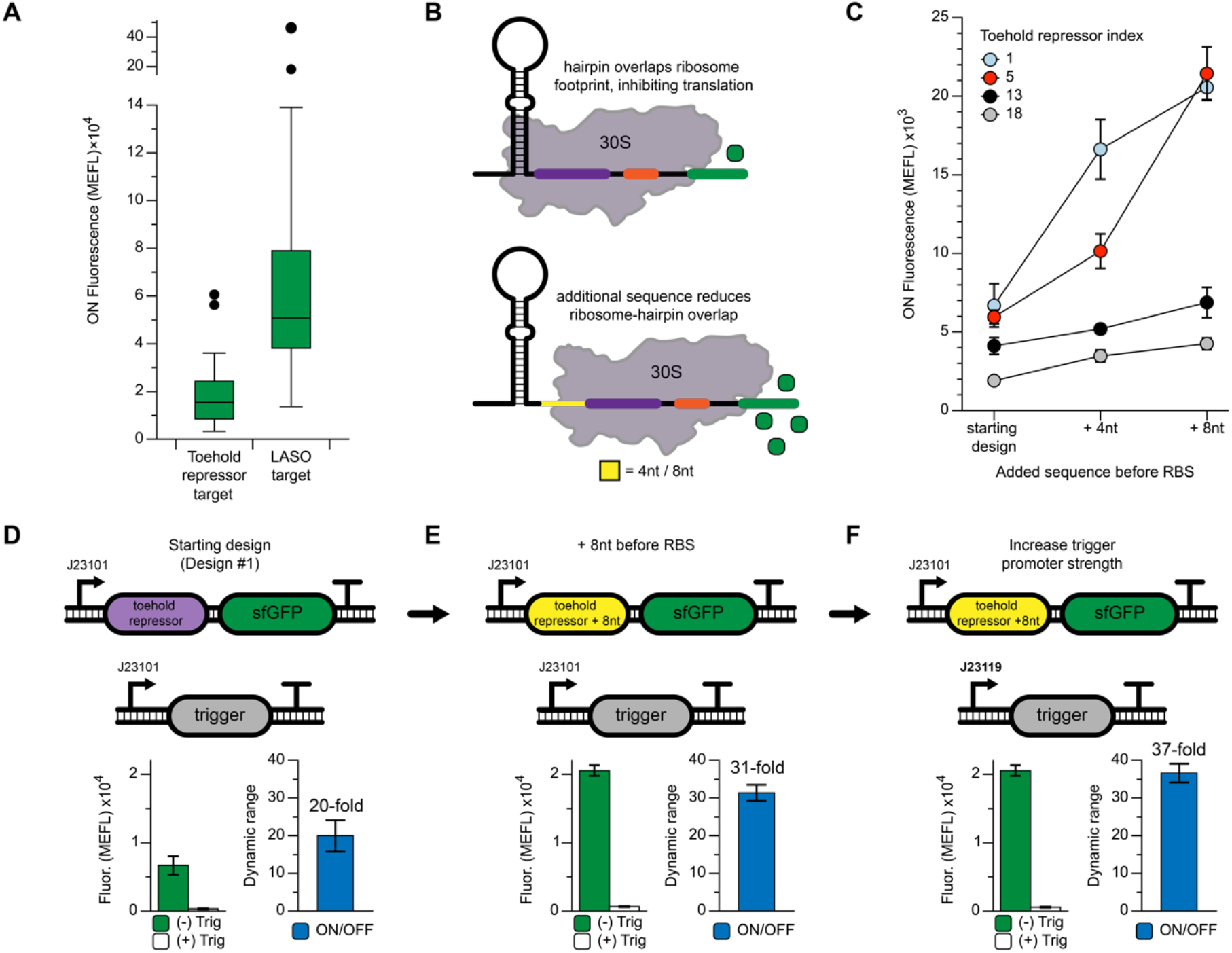
Strategies to improve the dynamic range of a toehold repressor. (A) Comparison of the ON (no trigger) fluorescence for the library of toehold repressors (from Figure 1C) and LASO designs (from Figure 1E). Despite identical RBS sequences across all designs, toehold repressors show a lower median ON level. (B) Proposed cause of reduced ON level of the toehold repressor design. Overlap between the ribosomal footprint and ON hairpin may inhibit translation initiation. Increasing the distance between the RBS and ON hairpin could reduce this inhibition. (C) Adding 4nt or 8nt of designed sequence before the RBS increases the ON level in four different toehold repressor designs. (D) Toehold repressor design #1 has a dynamic range (ON fluorescence divided by OFF fluorescence) of 20-fold. Adding 8nt before the RBS (E) increases the dynamic range to 31-fold mainly by increasing the ON level. (F) Increasing trigger expression by using a stronger promoter reduces the OFF level, which increases the dynamic range even further to 37-fold.

Previous studies have identified other strategies to increase the dynamic range of RNA regulators, including tuning the promoter strengths of target and trigger species^20^. The expression context used up until this point expresses both target and trigger RNAs using the same J23101 *E. coli* σ^70^ promoter^36^, but under different plasmid copy numbers to establish an excess of trigger RNA in the cell. To further increase this excess, the strength of the trigger promoter can be increased and/or the strength of the target promoter can be decreased. Using the improved design #1 (Figure 3E) as a test case, we cloned a stronger σ^70^ promoter (J23119) before the trigger, and a weaker promoter (J23150) before the target^43^. We then tested all promoter combinations to identify if altering the expression of one or both RNAs increased the dynamic range of the design (Supplementary Figure 7A). Strengthening the trigger promoter improved the dynamic range to 37-fold (Figure 3F), while weakening the target plasmid promoter did not improve performance (Supplementary Figure 7B).

We next tested the effect of altering the reporter protein sequence on the observed dynamic range. Previous work has found that degradation-tagged reporters can increase the dynamic range of a regulator by decreasing leak in the OFF state^33^. In the case of our optimized design #1, we observed a small increase in dynamic range to 41-fold using GFPmut3b-ASV, a degradation-tagged GFP variant. This increase was not significant compared to sfGFP, perhaps owing to the very low OFF level for this particular design even when using an sfGFP reporter (Supplementary Figure 7C,D).

Overall, these results illustrate an important advantage of *de novo*-designed riboregulators like toehold repressors. The large design space of these regulators allows many design variants to be tested, including changes to sequence, expression context, and lengths of subdomains like toeholds and hairpins. This flexibility allows the design to be extensively optimized to maximize the performance of the regulator.

### LASO designs can distinguish between closely-related transcripts

Having shown that our translational repressor designs can form orthogonal libraries using mRNA targets with high sequence diversity (Figure 2), we wanted to determine the limits of our LASO design in discriminating between closely-related transcripts. As a design principle, LASOs include specifically looped out regions of the target that are conserved. We hypothesized that avoiding direct base-pairing to these conserved sequences reduces the likelihood of crosstalk between designs. Given that translation start sites in *E. coli* are purine-rich and that the AUG start codon is used in ~83% of genes^38,39^, this approach may also help prevent unintended repression of endogenous mRNAs. As a test case, we used an MG1655-derived strain that expresses both sfGFP and mRFP from the same genomic locus^44^. This strain is a useful worst-case scenario for potential off-target repression, as the sfGFP and mRFP mRNAs have identical 5’ UTRs. To further increase the likelihood of crosstalk, we over-expressed LASO species using a strong *E. coli* σ^70^ promoter (J23119) and a high copy number plasmid (ColE1) to promote saturation of LASO abundance in the cell. To test designs, individual colonies were picked, grown overnight in LB, diluted 1:50 in M9 minimal media and grown for 4h. Absorbance (OD_600_), mRFP fluorescence, and sfGFP fluorescence were measured using a BioTek Synergy plate reader (see Materials and Methods).

Using mRFP as our desired target mRNA, we compared repression from a typical asRNA design - which forms a continuous duplex across the translation start site - to our LASO design that loops out the RBS and start codon in the binding interaction. As expected, the asRNA strongly repressed mRFP, but also showed high off-target sfGFP repression (77%), presumably due to base-pairing across the sfGFP 5’ UTR (Figure 4A). Interestingly, our LASO design maintained strong repression of mRFP, but off-target repression of sfGFP was limited to just 10% (Figure 4B). This result demonstrates the ability of our design motif to specifically target a gene of interest without repressing highly similar mRNAs.

**Figure 4.**
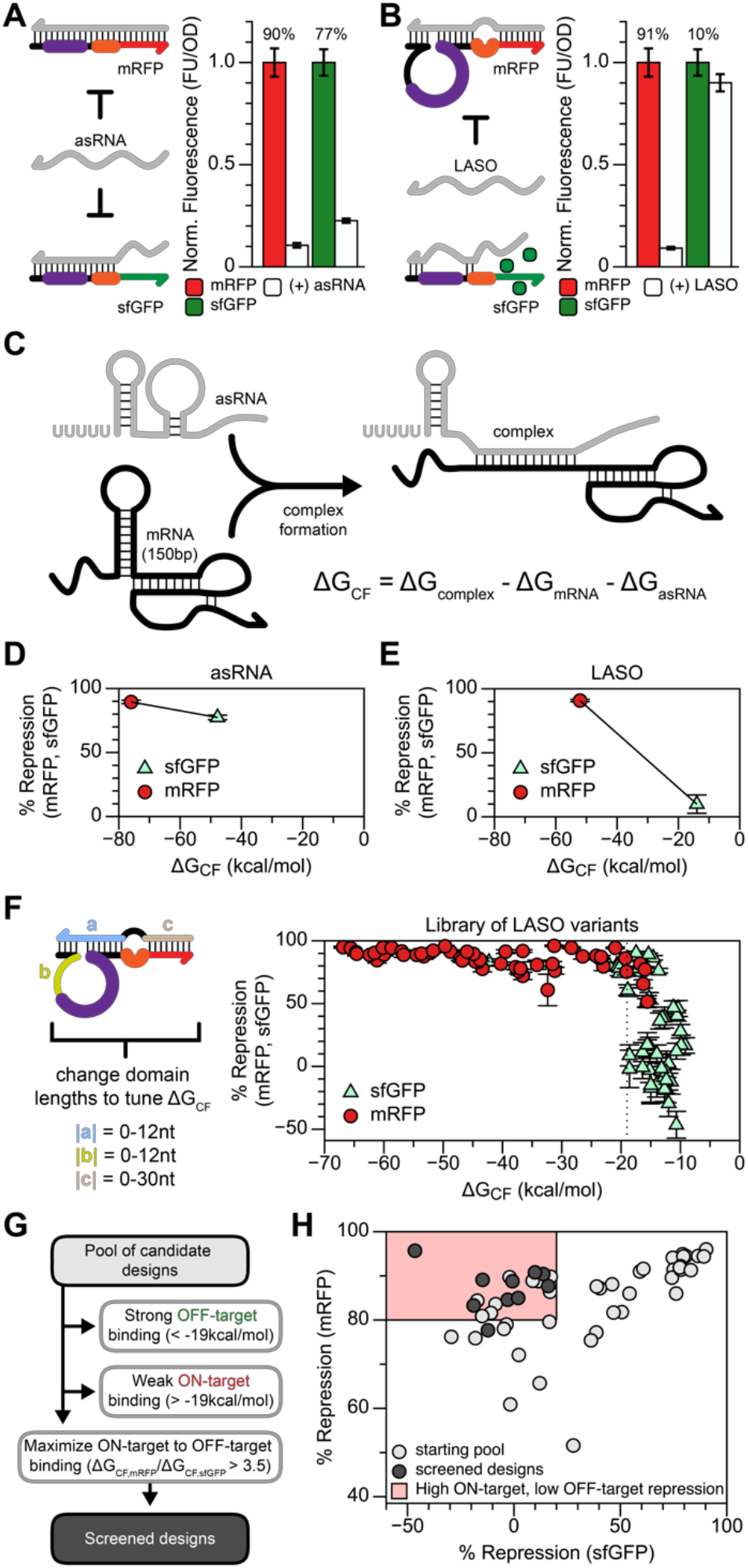
The LASO design enables strong repression of a chromosomally-encoded target gene without repressing a gene with a closely related 5’ UTR. (A) An MG1655-derived strain^44^ expresses both mRFP and sfGFP from a genomic locus, each using the same 5’ UTR. A typical asRNA design to inhibit mRFP translation binds across the translation start site and N-terminal coding sequence, repressing mRFP. However, sequence similarities cause unintended off-target repression of sfGFP. (B) The LASO design, which is designed to not directly bind the RBS or start codon of the shared 5’ UTR, shows robust on-target repression (mRFP) and low off-target repression (sfGFP). Percent repression indicated above conditions in (A,B). (C) The interaction between a mRNA and an asRNA can be modelled by estimating the free energy of complex formation (ΔG_CF_), which is the difference between the free energy of the complex minus the free energies of the isolated mRNA and asRNA structural states. (D) A low (stable) ΔG_CF_ < –40 kcal/mol is observed for both mRFP and sfGFP repression by the full asRNA design. (E) The LASO design shows a distinct correlation between reducing off target repression of sfGFP and a destabilized (higher) ΔG_CF_. (F) A library of LASOs targeting mRFP were constructed to tune the strength of interaction with the target, and ΔG_CF_ was calculated for both the mRFP and sfGFP expression cassettes. Comparing all on-target (mRFP) and off-target (sfGFP) repression to ΔG_CF_ shows a threshold of approximately –19 kcal/mol (indicated by a dashed line), below which designs more strongly repress translation and above which designs in aggregate show reduced cross talk. (G) A proposed workflow for screening optimal LASO designs. First, designs with strong predicted off-target binding (ΔG_CF,sfGFP_ < –19kcal/mol) and weak on-target binding (ΔG_CF,mRFP_ > –19kal/mol) are removed. Remaining designs with the highest ratio of on-target ΔG_CF_ to off-target ΔG_CF_ (ΔG_CF,RFP_/ΔG_CF,sfGFP_ > 3.5) are kept. (H) Applying these criteria to the starting pool of 52 LASO designs, 10 designs can be correctly identified all with low off-target (sfGFP) repression and at least 78% on-target (mRFP) repression.

We next sought to develop a computational strategy to predict on- and off-target repression effects, which could be incorporated into the design of LASOs. Na et al.^17^ showed that the calculated binding energy between an sRNA and mRNA (the ΔG of the duplex) can help explain differences in repression efficiency between length variants of an sRNA repressor. Hoynes-O’Connor and Moon^19^ similarly proposed using the free energy of complex formation between an asRNA and mRNA to predict asRNA performance. While this metric proved useful, it does not account for RNA structures within mRNA targets or elements of the asRNA outside of the binding region that compete with productive asRNA-mRNA binding. To remedy this, we updated this metric to calculate the free energy of complex formation (ΔG_CF_), defined as the free energy difference between the bound state, consisting of the productive asRNA-mRNA interaction (ΔG_complex_), and the unbound state, consisting of the sum of the folding free energies of the 5’ end of the mRNA (ΔG_mRNA_,150bp total) and the entire asRNA (ΔG_asRNA_), including the terminator and any sequence outside of the binding region (Figure 4C). We calculated ΔG_CF_ for the asRNA and LASO designs, and found that the asRNA showed low ΔG_CF_ values (< –40kcal/mol) for both mRFP and sfGFP (Figure 4D). Low ΔG_CF_ was also observed for LASO targeting to mRFP, but low off-target sfGFP repression corresponded to higher ΔG_CF_ (Figure 4E). This provided evidence that ΔG_CF_ could be a useful metric to minimize off-target repression.

To more rigorously investigate ΔG_CF_ as a LASO design parameter, we built a library of 52 LASO designs in which we changed the lengths of specific binding interactions to tune ΔG_CF_ (Figure 4F). We then measured repression of both mRFP and sfGFP from these LASO designs and found a stark pattern: below a ΔG_CF_ of approximately –19 kcal/mol there is a trend of increasing repression with decreasing ΔG_CF_, which plateaus at very low values of ΔG_CF_; above a ΔG_CF_ of approximately –19 kcal/mol there is a range of observed repression (Figure 4F). This general trend is consistent with previous results^19^; however, incorporating the structure of the mRNA target into the calculation of ΔG_CF_ allowed for a more clear and direct comparison between genes.

We next sought to develop a set of design criteria to screen candidate LASO designs to identify variants with high on-target repression (> 80% repression of mRFP) and weak off-target repression (< 20% repression of sfGFP). Using calculations of ΔG_CF_, we developed a set of 3 design criteria that identify LASO designs with these properties from a larger starting pool of designs (Figure 4G). First, designs with strong predicted off-target binding (ΔG_CF_ < –19kcal/mol) are removed. Second, designs with weak on-target binding (ΔG_CF_ > –19kal/mol) are removed. Finally, designs with a high, empirically-determined ratio of on-target to off-target ΔG_CF_ (ΔG_CF,mRFP_/ΔG_CF,sfGFP_ > 3.5) are kept. Applying these criteria to the starting pool of 52 LASO designs, we could correctly identify 10 designs, all with low off-target sfGFP repression (Figure 4H, Supplementary Figures 8 and 9). Additionally, every design except one showed high mRFP repression; the one outlier gave just below 80% repression (78% ± 2.4%). These results show how simple *in silico* calculations can be used to identify LASO designs that can discriminate between closely-related transcripts.

### LASO designs can predictably repress endogenous mRNA sequences

An important application of asRNAs and sRNAs is translational repression of endogenous genes in *E. coli*. To demonstrate the ability of LASOs to repress sequences derived from natively expressed mRNAs, we created fusion proteins where the 5’ UTR and initial coding region of several *E. coli* genes were fused to sfGFP (Figure 5A). LASOs were designed to target the translation start site of these fusion constructs (Figure 5B), and the repression performance was measured by flow cytometry (see Materials and Methods). Significant repression was observed in each case (Figure 5C), although the strength of repression ranged from 42% to 94%. We hypothesized that structural inaccessibility of the mRNA target may inhibit mRNA/LASO complex formation and cause this range of observed repression. As expected based on our previous study of mRFP and sfGFP targeting, repression strength decreased with increasing ΔG_CF_ (Figure 5D). This suggests that our updated definition of ΔG_CF_ is a useful heuristic to not just to capture off-target interactions, but to predict the strength of repression of mRNA targets with varying degrees of structural inaccessibility (Supplementary Figure 10).

**Figure 5.**
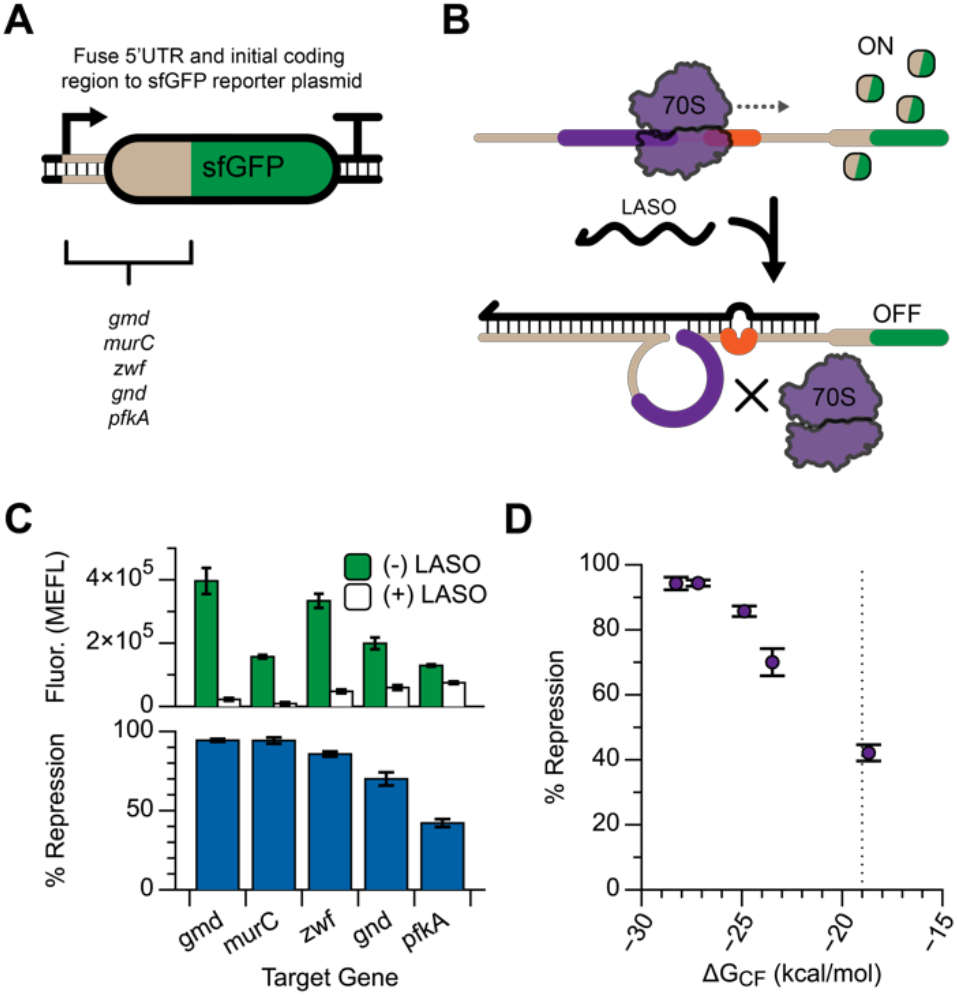
The LASO design can predictably target mRNA sequences from endogenous genes. The 5’ UTR and initial coding region (51nt corresponding to 17AA) of several endogenous *E. coli* genes were translationally fused to an sfGFP coding sequence to create a chimeric expression cassette (A), and LASOs were designed to target the expressed chimeric mRNAs (B). (C) Each target was responsive to LASO expression, with repression efficiency ranging from 94% to 42%. (D) Differences in repression can be explained by calculated ΔG_CF_ values for each LASO/target pair, with stronger repression corresponding to lower values of ΔG_CF_ (−19 kcal/mol indicated by a dashed line).

### LASO designs enable one-step tuning of RBS strength

Applications in metabolic engineering and synthetic biology often require iterative cycles of tuning gene expression levels, which is often accomplished by modifying the strength of translation initiation^18,45-47^. This is typically done by changing the sequence of the RBS and/or start codon, either by random mutagenesis or forward design. These sequence changes require compensatory mutations to any asRNA designed to bind to the translation start site, which is disadvantageous when troubleshooting individual components of a larger gene expression circuit. In many cases, the entire circuit must be re-designed to compensate for the sequence changes; at a minimum, libraries of asRNA variants must be re-tested when the RBS sequence of their target is changed. Conveniently, the LASO design of looping out the translation initiation region of the mRNA from the binding interaction provides a simple method of tuning RBS strength without requiring sequence changes to the asRNA. As a proof-of-principle, we introduced a degenerate RBS sequence (NNNNNNNN) and start codon (RTG) into LASO target #27 by iPCR (Figure 6A). Colonies of varying sfGFP expression strength were picked by eye using a blue light tray, and the performance of each library member was tested using the same LASO design. A range of ON levels spanning more than 2 orders of magnitude was observed. When characterized with LASO expression, repression exceeding 90% was observed over most of this range (Figure 6B), although some loss in repression efficiency was observed for library members with very low ON levels.

**Figure 6.**
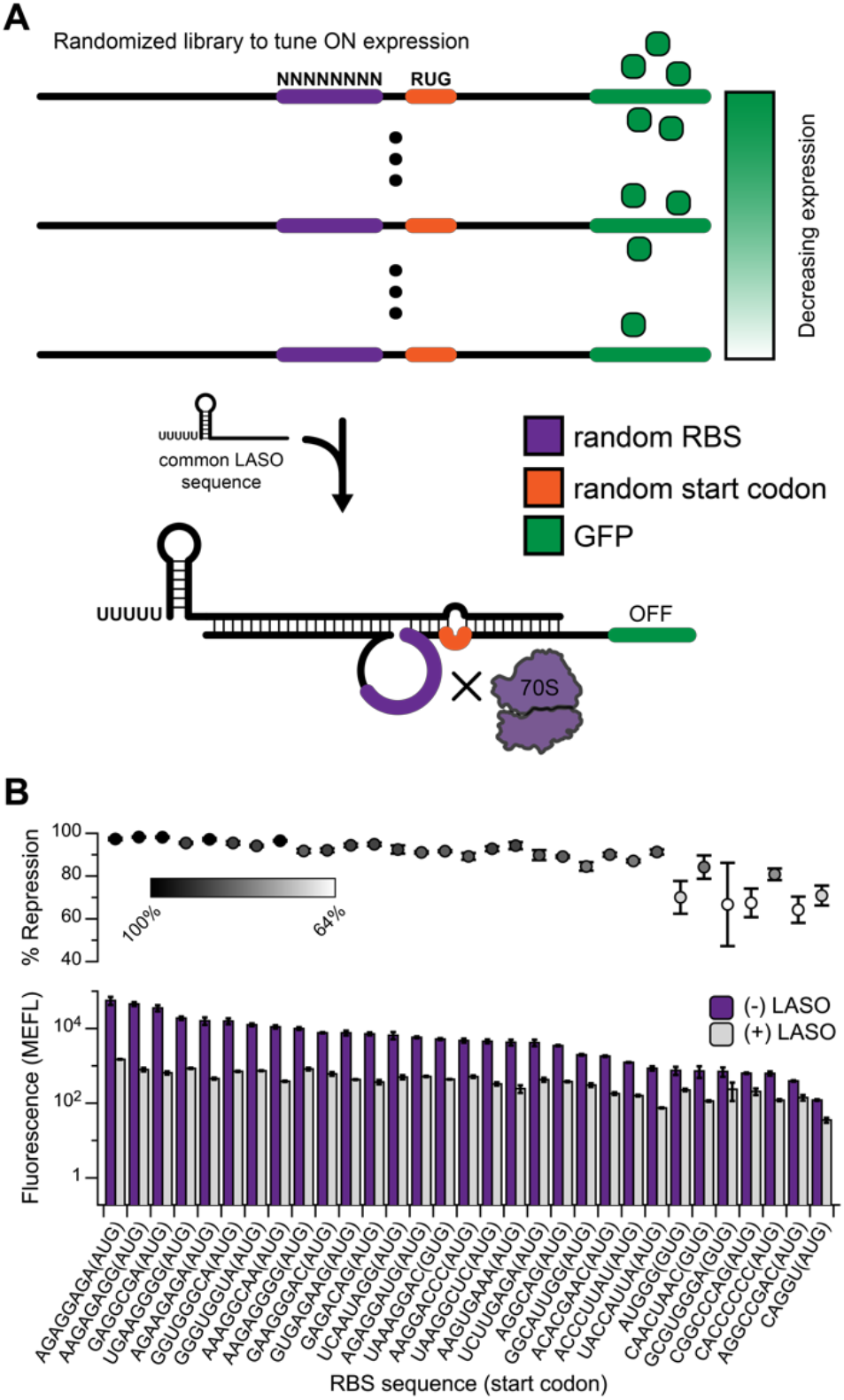
The LASO design enables facile tuning of RBS strength. (A) The expression of a target mRNA can be tuned in a single step by the introduction of a degenerate RBS and start codon, generating a library with a range of ON levels. A single LASO can be used to repress all library members because it avoids direct binding to the mutated sequences. Using this method, 31 functional target constructs were generated in a single step (B), with ON fluorescence spanning more than 2 orders of magnitude, with 24 designs showing greater than 85% repression.

We chose to tune expression by screening random sequences, but tools such as the RBS Calculator^42,46^ can be used to design RBS sequences if the desired translation rate is known. We found that the ON levels of the library members generated by random mutagenesis were consistent with predicted translation rates using RBS Calculator v2.0 (Supplementary Figure 11), providing support that either approach (screening or forward design) could be used for simple, one-step expression tuning of target/LASO pairs.

### Toehold regulators enable simultaneous activation and repression of unrelated genes from a single input RNA

Finally, we wanted to investigate whether the translational RNA repressors developed in this work are functionally compatible with other RNA regulators. Specifically, we sought to verify that two different riboregulators can function simultaneously in the same cell to independently and differently regulate two different genes. We designed a toehold switch (controlling mRFP) and a toehold repressor (controlling sfGFP) to respond to the same trigger RNA sequence (Figure 7A). We confirmed that each regulator functioned as expected when tested in isolation (Figure 7B), and then tested both regulators in the same cell. Repression and activation performance were retained when the two regulators were co-expressed (Figure 7C), and switching was visible by eye (Supplementary Figure 12). This combined toehold switch-toehold repressor allowed the cells to be switched from expressing sfGFP (no trigger) to mRFP (with trigger), demonstrating the simultaneous activation and repression of two unrelated genes from a single ncRNA input.

**Figure 7.**
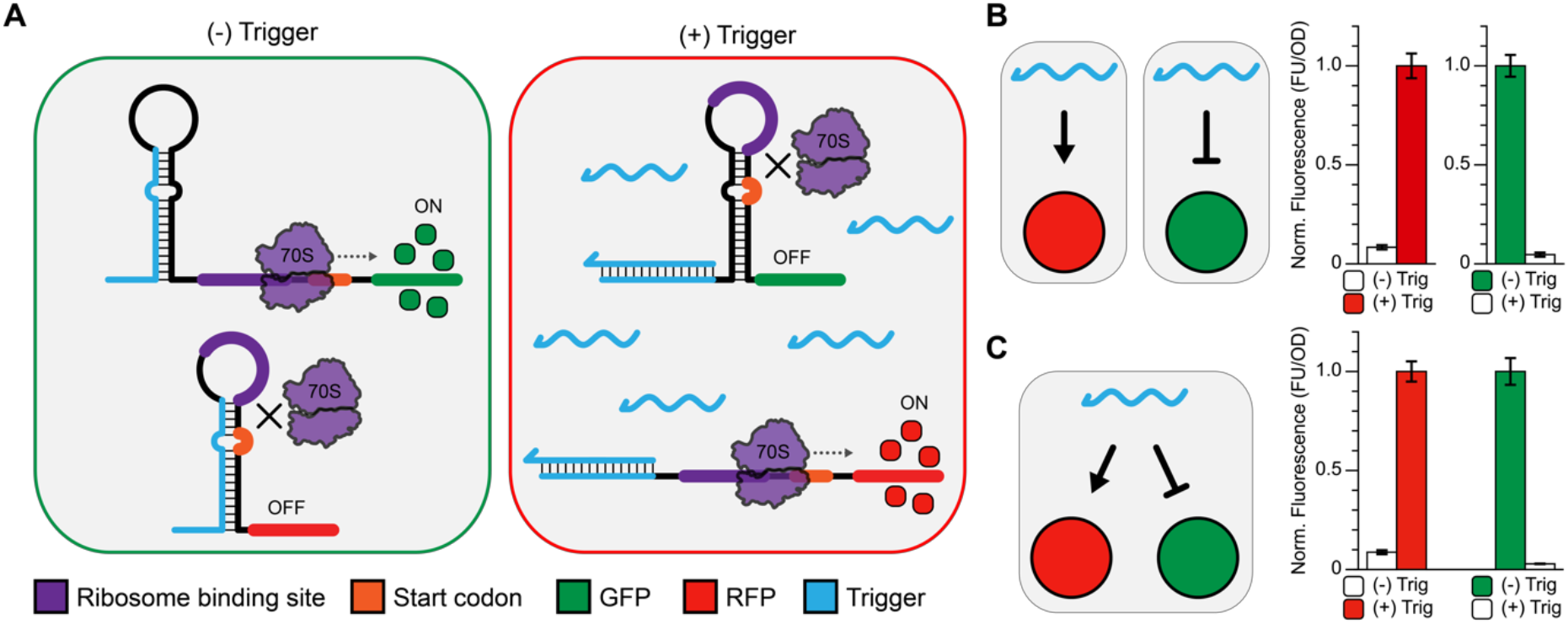
Simultaneous activation and repression of two genes from a single RNA input. (A) A toehold switch (controlling mRFP) and a toehold repressor (controlling sfGFP) were designed to interact with the same trigger RNA. When the trigger is not expressed, only the toehold repressor is in the ON state, resulting in sfGFP production in the cell. Trigger expression activates the toehold switch (producing mRFP) while simultaneously inhibiting sfGFP expression from the toehold repressor. (B) Performance of the toehold switch and toehold repressor when tested individually. (C) Performance of the toehold switch and repressor are maintained when co-expressed in the same cell.

## DISCUSSION

In this work, we have developed two strategies to create translational RNA repressors: toehold repressors and looped antisense oligonucleotides (LASOs) to directly repress translation by interacting with mRNA targets. Using rational RNA engineering strategies, we then successfully improved the performance of our toehold repressor design. These efforts enabled us to achieve some of the largest dynamic ranges reported for RNA-only repressors and identify sets of mutually orthogonal target/trigger pairs, serving as an important step forward for the practical implementation of RNA-based networks to control gene expression.

Toehold repressors fill an important gap in the available toolbox of riboregulators. Like toehold switches, the toehold repressor can be designed *in silico*, without the sequence constraints of previously-reported design strategies^41^. This design flexibility, as well as structural and mechanistic similarities between toehold switches and toehold repressors, makes integration of activation and repression into the same genetic network a straightforward task (Figure 7). This is a marked improvement over trying to integrate multiple regulatory mechanisms that are currently available, which is difficult due to sequence constraints on input and/or output RNA species. The similar structures of the toehold switch and the ON hairpin of the toehold repressor (Figure 1A,B) also create the possibility of inter-converting toehold switches to repressors (and vice-versa), which would aid in the refactoring of genetic network components without altering the sequence of their corresponding trigger RNAs.

Perhaps more important is our development of the LASO design concept for translational repression. By merging the concept of asRNA inhibition through asRNA-mRNA interactions with the design features of toehold switches and repressors – namely eliminating base-pairing to the RBS and start codon by creating looped regions of the binding interaction – we have developed a new concept for antisense-RNA mediated translational repression that is highly efficient, predictable and modular. Specifically, we demonstrated that our LASO design can strongly repress target mRNAs without crosstalk with closely-related transcripts and without the inclusion of an Hfq-binding scaffold common to other sRNA translational repressor designs. This design strategy also allows for single-step tuning of RBS strength without changing the sequence of the LASO, enabling facile screening or forward-design of repressible expression constructs with desired ON expression levels.

We also demonstrated that LASO designs can be computationally assessed for design quality. Specifically, a thorough characterization of off-target interactions in a dual-reporter assay (Figure 4) led us to propose an improved metric for predicting asRNA-mRNA interactions. By adding terms that account for competing structures to the measurement of the free energy of complex formation (ΔG_CF_), we showed that this metric can be used to capture off-target interactions as well as inefficient target repression arising from structural inhibition of the target mRNA. This is a straightforward and computationally inexpensive calculation which could be routinely incorporated into the forward design of new sRNA repressors, facilitating the *in silico* screening of hundreds to thousands of potential off-target interactions for a single design. Furthermore, these simple design principles, combined with the ability of LASOs to repress targets without any modification of the mRNA sequence, could enable application of this concept to a broad array of work requiring modulation of endogenous gene expression.

While the design rules developed in this work are directly relevant to applications including protein expression balancing or screening gene knockdowns for metabolic pathways engineering^17,18^, they are also important for asRNA design more broadly. Antisense oligonucleotides have emerged as a promising class of therapeutic drugs that correct errors in gene expression through asRNA-mRNA interactions^35,48^. Their utility will only expand as new cases of RNA-based genetic misregulation are uncovered^49,50^. Balancing target site accessibility with potential off-target binding is critical for the safety and efficacy of nucleic acid-based therapeutics. The design flexibility of the LASO concept – where specific regions can be looped out of the productive binding interaction – could offer great benefits when designing antisense oligonucleotide therapies. For example, for complex targets that have structures with only small windows available for binding, asRNAs that are designed to bind to full contiguous regions may fail through the formation of intramolecular structures within the asRNA. However, LASOs designed and screened to bind specifically to the available regions may function. Future work in this area could also benefit greatly from the use of RNA structure probing^50-53^ to identify the structurally accessible regions of complex targets, and refine calculations of ΔG_CF_^51^. This would enable more accurate characterization of target site accessibility, especially in the cellular context where factors like RNA-protein interactions can confound predictions of RNA accessibility. It is likely that the LASO design motifs, applied to translational control in this work, will also be useful in other contexts where RNA-RNA interactions can modulate gene expression.

While this work was being performed, an independent study identified similar strategies for creating RNA repressors (Kim et al., Submitted). In addition to the toehold repressor strategy identified here, they identified a third repressor design which relies on the conditional formation of a stabilized three-way junction (3WJ) structure to enable up to 4-input logic computation. Together, these studies underscore the power of *de novo* RNA design to address a range of previously unmet design challenges, ranging from biomolecular logic computation to the repression of endogenous genes with a high degree of predictability and specificity.

In summary, we have shown that translational RNA repressors can be designed *in silico* using first principles with no hard sequence constraints on the trigger RNA input. This work serves as a significant step forward in the development of a versatile toolbox of RNA-based gene expression regulators for synthetic biology and metabolic engineering, and provides a framework for the refinement of antisense oligonucleotide design more broadly.

## Methods

### Cloning and plasmid construction

Toehold repressor sequences were synthesized as single-stranded fragments (IDT DNA), amplified using common flanking primers, and introduced by Gibson Assembly into a p15A plasmid backbone harbouring chloramphenicol resistance. All other sequences were constructed by inverse PCR (iPCR). Trigger/LASO sequences were cloned into ColE1 backbones harbouring carbenecillin resistance. The toehold switch tested in Figure 7 was cloned into a CDF backbone harbouring spectinomycin resistance. See supplemental information for plasmid architectures (Supplementary Figure 2) and key sequences (Supplementary Tables 1 and 2).

### Flow cytometry data collection

All flow cytometry experiments were performed in *E. coli* strain TG1 (*F’traD36 lacIq Delta(lacZ) M15 pro A+B+/supE Delta(hsdM-mcrB)5 (rk- mk- McrB-) thi Delta(lac-proAB)*). Cells were transformed with the relevant combination of target and trigger/LASO plasmids or target plasmid with pJBL002 (a blank trigger plasmid). An autofluorescence control was included by transforming blank target and trigger plasmids (pJBL001 and pJBL002, respectively) which were both taken from Lucks et al.^37^. Plasmid combinations were transformed into chemically competent *E. coli* TG1 cells, plated on Difco LB+Agar plates containing 100 μg/mL carbenicillin and 34 μg/mL chloramphenicol, and grown overnight at 37 °C. Following overnight incubation, plates were left at room temperature for approximately 9 h. Individual colonies were grown overnight in LB, then diluted 1:50 into M9 minimal media. After 6 h, cells were diluted 1:100 in 1× Phosphate Buffered Saline (PBS) containing 2 mg/mL kanamycin. A BD Accuri C6 Plus flow cytometer fitted with a high-throughput sampler was then used to measure sfGFP fluorescence. Measurements were taken for at least 6 biological replicates collected over at least two days, with one exception; owing to the large number of transformations, data presented in Figure 2 represents at least 3 biological replicates collected on a single day.

### Flow cytometry data analysis

Flow cytometry data analysis was performed using FlowJo (v10.4.1). Cells were gated by FSC-A and SSC-A, and the same gate was used for all samples. Samples exhibiting non-unimodal fluorescence distributions were excluded from further analysis, and the geometric mean fluorescence was calculated for each sample. All fluorescence measurements were converted to Molecules of Equivalent Fluorescein (MEFL) using CS&T RUO Beads (BD cat#661414), which were run on each day of data collection. The average fluorescence (MEFL) over replicates of cells expressing empty plasmids (pJBL001 and pJBL002)^37^ was then subtracted from each measured fluorescence value.

### *In vivo* bulk fluorescence data collection

Bulk fluorescence experiments were performed in *E. coli* strain TG1 (*F’traD36 lacIq Delta(lacZ) M15 pro A+B+/supE Delta(hsdM-mcrB)5 (rk- mk- McrB-) thi Delta(lac-proAB)*), MG1655 (*F^−^ λ^−^ ilvG^−^ rfb-50 rph-1*), or an MG1655-derived strain expressing mRFP and sfGFP^44^. Designs targeting mRFP were transformed with a trigger plasmid only, because the target was expressed from the genome; in this case, the parent strain (MG1655) transformed with pJBL002 served as the autofluorescence control. Transformed cells were plated on Difco LB+Agar plates containing 100 μg/mL carbenicillin and grown overnight at 37 °C. Following overnight incubation on plates, individual colonies were grown overnight in LB, then diluted 1:50 into M9 minimal media. After 4 h, sfGFP fluorescence, mRFP fluorescence, and optical density (OD_600_) were measured using a Biotek Synergy plate reader. Measurements were taken for at least 6 biological replicates collected over at least two days.

Testing of a toehold switch and toehold repressor in parallel (Figure 7 and Supplementary Figure 12) was performed using a 3-plasmid system, with trigger and toehold repressor target expressed as described previously (Supplementary Figure 2A,B). The toehold switch was expressed from a CDF-based plasmid harbouring spectinomycin resistance (Supplementary Figure 2C). The pCDF-Duet plasmid (Novagen) was used as the blank control plasmid for the toehold switch. Cells were plated on Difco LB+Agar plates containing 100 μg/mL carbenicillin, 34 μg/mL chloramphenicol, and 100 μg/mL spectinomycin. Following overnight incubation at 37 °C, individual colonies were grown overnight in LB, then diluted 1:50 into M9 minimal media. After 6 h, sfGFP fluorescence, mRFP fluorescence, and optical density (OD_600_) were measured using a Biotek Synergy plate reader. Measurements were taken for at least 6 biological replicates collected over at least two days.

### Bulk fluorescence data analysis

Bulk florescence data for sfGFP and mRFP were analysed in a similar manner to the flow cytometry experiments, except that all fluorescence values were normalized to absorbance (OD_600_). First, the OD_600_ of a blank well (containing only M9 media) was subtracted from the OD_600_ of each well. Then, FL/OD_600_ was calculated for each well containing cells; the average FL/OD_600_ of the autofluorescence control was then subtracted from each experimental well to determine the final, normalized value of FL/OD_600_ for each condition. Following calculations of average and standard deviation for each experimental condition, data were re-normalized such that the ON fluorescence of sfGFP and mRFP was set to 1 (Figure 4A,B and Figure 7B,C).

## ACKNOWLEDGEMENTS

The authors thank Melissa Takahashi and Elizabeth Weiss for assistance with preliminary data collection.

## FUNDING

This work was supported by the National Science Foundation Graduate Research Fellowship Program [grant number DGE-1144153 to PDC and CJG], an NSF CAREER award (1452441 to J. B. L.), and Searle Funds at The Chicago Community Trust (to J. B. L.).

## AUTHOR CONTRIBUTIONS

P.D.C. and J.B.L. conceived and designed this study, and P.D.C. and C.J.G. performed experiments.

## COMPETING INTERESTS

The authors declare no competing financial interests.

