## Supplementary Information for "De novo Design of Translational RNA Repressors"

Paul D. Carlson[1,2], Cameron J. Glasscock[1,2], and Julius B. Lucks[2,3,*]
[1] Robert F. Smith School of Chemical and Biomolecular Engineering, Cornell University, Ithaca, NY, 14853, USA
[2] Center for Synthetic Biology, Northwestern University, Evanston, IL, 60208, USA
[3] Department of Chemical and Biological Engineering, Northwestern University, Evanston, IL, 60208, USA

**Supplementary Table 1.** Important sequences used in this study. Plasmid architectures shown in Supplementary Figure 1 and Supplementary Table 2.

| Name | Sequence |
|---|---|
| J23101 promoter | TTTACAGCTAGCTCAGTCCTAGGTATTATGCTAGC |
| J23119 promoter | TTGACAGCTAGCTCAGTCCTAGGTATAATAGATC |
| J23119 promoter (with SpeI cut site) | TTGACAGCTAGCTCAGTCCTAGGTATAATACTAGT |
| J23150 promoter | TTTACGGCTAGCTCAGTCCTAGGTATTATGCTAGC |
| Superfolder GFP (sfGFP) | ATGAGCAAAGGAGAAGAACTTTTCACTGGAGTTGTCCCAAT TCTTGTTGAATTAGATGGTGATGTTAATGGGCACAAATTTTC TGTCCGTGGAGAGGGTGAAGGTGATGCTACAAACGGAAAA CTCACCCTTAAATTTATTTGCACTACTGGAAAACTACCTGTT CCGTGGCCAACACTTGTCACTACTCTGACCTATGGTGTTCA ATGCTTTTCCCGTTATCCGGATCACATGAAACGGCATGACT TTTTCAAGAGTGCCATGCCCGAAGGTTATGTACAGGAACGC ACTATATCTTTCAAAGATGACGGGACCTACAAGACGCGTGC TGAAGTCAAGTTTGAAGGTGATACCCTTGTTAATCGTATCGA GTTAAAGGGTATTGATTTTAAAGAAGATGGAAACATTCTTGG ACACAAACTCGAGTACAACTTTAACTCACACAATGTATACAT CACGGCAGACAAACAAAAGAATGGAATCAAAGCTAACTTCA AAATTCGCCACAACGTTGAAGATGGTTCCGTTCAACTAGCA GACCATTATCAACAAAATACTCCAATTGGCGATGGCCCTGT CCTTTTACCAGACAACCATTACCTGTCGACACAATCTGTCCT TTCGAAAGATCCCAACGAAAAGCGTGACCACATGGTCCTTC TTGAGTTTGTAACTGCTGCTGGGATTACACATGGCATGGAT GAGCTCTACAAATAA |
| GFPmu3b-ASV | ATGCGTAAAGGAGAAGAACTTTTCACTGGAGTTGTCCCAAT TCTTGTTGAATTAGATGGTGATGTTAATGGGCACAAATTTTC TGTCAGTGGAGAGGGTGAAGGTGATGCAACATACGGAAAA CTTACCCTTAAATTTATTTGCACTACTGGAAAACTACCTGTT CCGTGGCCAACACTTGTCACTACTTTCGGTTATGGTGTTCA ATGCTTTGCGAGATACCCAGATCACATGAAACAGCATGACT |

| | |
|---|---|
| | TTTTCAAGAGTGCCATGCCCGAAGGTTACGTACAGGAAAGA ACTATATTTTTCAAAGATGACGGGAACTACAAGACACGTGC TGAAGTCAAGTTTGAAGGTGATACCCTTGTTAATAGAATCGA GTTAAAAGGTATTGATTTTAAAGAAGATGGAAACATTCTTGG ACACAAATTGGAATACAACTATAACTCACACAATGTATACAT CATGGCAGACAAACAAAAGAATGGAATCAAAGTTAACTTCA AAATTAGACACAACATTGAAGATGGAAGCGTTCAACTAGCA GACCATTATCAACAAAATACTCCGATTGGCGATGGCCCTGT CCTTTTACCAGACAACCATTACCTGTCCACACAATCTGCCCT TTCGAAAGATCCCAACGAAAAGAGAGACCACATGGTCCTTC TTGAGTTTGTAACCGCTGCTGGGATTACACATGGCATGGAT GAACTATACAAAAGGCCTGCAGCAAACGACGAAAACTACGC TGCATCAGTTTAATAA |
| Monomeric red fluorescent protein (mRFP) | ATGGCGAGTAGCGAAGACGTTATCAAAGAGTTCATGCGTTT CAAAGTTCGTATGGAAGGTTCCGTTAACGGTCACGAGTTCG AAATCGAAGGTGAAGGTGAAGGTCGTCCGTACGAAGGTAC CCAGACCGCTAAACTGAAAGTTACCAAAGGTGGTCCGCTG CCGTTCGCTTGGGACATCCTGTCCCCGCAGTTCCAGTACG GTTCCAAAGCTTACGTTAAACACCCGGCTGACATCCCGGAC TACCTGAAACTGTCCTTCCCGGAAGGTTTCAAATGGGAACG TGTTATGAACTTCGAAGACGGTGGTGTTGTTACCGTTACCC AGGACTCCTCCCTGCAAGACGGTGAGTTCATCTACAAAGTT AAACTGCGTGGTACCAACTTCCCGTCCGACGGTCCGGTTAT GCAGAAAAAAACCATGGGTTGGGAAGCTTCCACCGAACGT ATGTACCCGGAAGACGGTGCTCTGAAAGGTGAAATCAAAAT GCGTCTGAAACTGAAAGACGGTGGTCACTACGACGCTGAA GTTAAAACCACCTACATGGCTAAAAAACCGGTTCAGCTGCC GGGTGCTTACAAAACCGACATCAAACTGGACATCACCTCCC ACAACGAAGACTACACCATCGTTGAACAGTACGAACGTGCT GAAGGTCGTCACTCCACCGGTGCTTAA |
| TrrnB terminator (BamHI-BglII scar) | GGATCTGAAGCTTGGGCCCGAACAAAAACTCATCTCAGAAG AGGATCTGAATAGCGCCGTCGACCATCATCATCATCATCAT TGAGTTTAAACGGTCTCCAGCTTGGCTGTTTTGGCGGATGA GAGAAGATTTTCAGCCTGATACAGATTAAATCAGAACGCAG AAGCGGTCTGATAAAACAGAATTTGCCTGGCGGCAGTAGC GCGGTGGTCCCACCTGACCCCATGCCGAACTCAGAAGTGA AACGCCGTAGCGCCGATGGTAGTGTGGGGTCTCCCCATGC GAGAGTAGGGAACTGCCAGGCATCAAATAAAACGAAAGGC TCAGTCGAAAGACTGGGCCTTTCGTTTTATCTGTTGTTTGTC GGTGAACT |
| T500 terminator (BamHI-BglII scar) | GGATCTCAAAGCCCGCCGAAAGGCGGGCTTTTTTTT |
| Double terminator | CCAGGCATCAAATAAAACGAAAGGCTCAGTCGAAAGACTGG GCCTTTCGTTTTATCTGTTGTTTGTCGGTGAACGCTCTCTAC |

| | |
|---|---|
| | TAGAGTCACACTGGCTCACCTTCGGGTGGGCCTTTCTGCGT<br>TTATA |
| cmR<br>(BamHI-BglII scar) | GGCACGTAAGAGGTTCCAACTTTCACCATAATGAAATAAGA<br>TCACTACCGGGCGTATTTTTTGAGTTATCGAGATTTTCAGGA<br>GCTAAGGAAGCTAAAATGGAGAAAAAAATCACTGGATATAC<br>CACCGTTGATATATCCCAATGGCATCGTAAAGAACATTTTGA<br>GGCATTTCAGTCAGTTGCTCAATGTACCTATAACCAGACCG<br>TTCAGCTGGATATTACGGCCTTTTTAAAGACCGTAAAGAAAA<br>ATAAGCACAAGTTTTATCCGGCCTTTATTCACATTCTTGCCC<br>GCCTGATGAATGCTCATCCGGAATTTCGTATGGCAATGAAA<br>GACGGTGAGCTGGTGATATGGGATAGTGTTCACCCTTGTTA<br>CACCGTTTTCCATGAGCAAACTGAAACGTTTTCATCGCTCT<br>GGAGTGAATACCACGACGATTTCCGGCAGTTTCTACACATA<br>TATTCGCAAGATGTGGCGTGTTACGGTGAAAACCTGGCCTA<br>TTTCCCTAAAGGGTTTATTGAGAATATGTTTTTCGTCTCAGC<br>CAATCCCTGGGTGAGTTTCACCAGTTTTGATTTAAACGTGG<br>CCAATATGGACAACTTCTTCGCCCCCGTTTTCACCATGGGC<br>AAATATTATACGCAAGGCGACAAGGTGCTGATGCCGCTGG<br>CGATTCAGGTTCATCATGCCGTTTGTGATGGCTTCCATGTC<br>GGCAGAATGCTTAATGAATTACAACAGTACTGCGATGAGTG<br>GCAGGGCGGGGCGTAATTTGATATCGAGCTCGCTTGGACT<br>CCTGTTGATAGATCCAGTAATGACCTCAGAACTCCATCTGG<br>ATTTGTTCAGAACGCTCGGTTGCCGCCGGGCGTTTTTTATT |
| ampR | CCAATGCTTAATCAGTGAGGCACCTATCTCAGCGATCTGTC<br>TATTTCGTTCATCCATAGTTGCCTGACTCCCCGTCGTGTAG<br>ATAACTACGATACGGGAGGGCTTACCATCTGGCCCCAGTG<br>CTGCAATGATACCGCGAGACCCACGCTCACCGGCTCCAGA<br>TTTATCAGCAATAAACCAGCCAGCCGGAAGGGCCGAGCGC<br>AGAAGTGGTCCTGCAACTTTATCCGCCTCCATCCAGTCTAT<br>TAATTGTTGCCGGGAAGCTAGAGTAAGTAGTTCGCCAGTTA<br>ATAGTTTGCGCAACGTTGTTGCCATTGCTACAGGCATCGTG<br>GTGTCACGCTCGTCGTTTGGTATGGCTTCATTCAGCTCCGG<br>TTCCCAACGATCAAGGCGAGTTACATGATCCCCCATGTTGT<br>GCAAAAAAGCGGTTAGCTCCTTCGGTCCTCCGATCGTTGTC<br>AGAAGTAAGTTGGCCGCAGTGTTATCACTCATGGTTATGGC<br>AGCACTGCATAATTCTCTTACTGTCATGCCATCCGTAAGATG<br>CTTTTCTGTGACTGGTGAGTACTCAACCAAGTCATTCTGAG<br>AATAGTGTATGCGGCGACCGAGTTGCTCTTGCCCGGCGTC<br>AATACGGGATAATACCGCGCCACATAGCAGAACTTTAAAAG<br>TGCTCAT |
| specR | TTATTTGCCGACTACCTTGGTGATCTCGCCTTTCACGTAGT<br>GGACAAATTCTTCCAACTGATCTGCGCGCGAGGCCAAGCG<br>ATCTTCTTCTTGTCCAAGATAAGCCTGTCTAGCTTCAAGTAT<br>GACGGGCTGATACTGGGCCGGCAGGCGCTCCATTGCCCA |

| | |
|---|---|
| | GTCGGCAGCGACATCCTTCGGCGCGATTTTGCCGGTTACT<br>GCGCTGTACCAAATGCGGGACAACGTAAGCACTACATTTCG<br>CTCATCGCCAGCCCAGTCGGGCGGCGAGTTCCATAGCGTT<br>AAGGTTTCATTTAGCGCCTCAAATAGATCCTGTTCAGGAAC<br>CGGATCAAAGAGTTCCTCCGCCGCTGGACCTACCAAGGCA<br>ACGCTATGTTCTCTTGCTTTTGTCAGCAAGATAGCCAGATCA<br>ATGTCGATCGTGGCTGGCTCGAAGATACCTGCAAGAATGTC<br>ATTGCGCTGCCATTCTCCAAATTGCAGTTCGCGCTTAGCTG<br>GATAACGCCACGGAATGATGTCGTCGTGCACAACAATGGT<br>GACTTCTACAGCGCGGAGAATCTCGCTCTCTCCAGGGGAA<br>GCCGAAGTTTCCAAAAGGTCGTTGATCAAAGCTCGCCGCGT<br>TGTTTCATCAAGCCTTACGGTCACCGTAACCAGCAAATCAA<br>TATCACTGTGTGGCTTCAGGCCGCCATCCACTGCGGAGCC<br>GTACAAATGTACGGCCAGCAACGTCGGTTCGAGATGGCGC<br>TCGATGACGCCAACTACCTCTGATAGTTGAGTCGATACTTC<br>GGCGATCACCGCTTCCCTCATACTCTTCCTTTTTCAATATTA<br>TTGAAGCATTTATCAGGGTTATTGTCTCATGAGCGGATACAT<br>ATTTGAATGTATTTAGAAAAATAAACAAA |
| p15A origin of replication | GCGCTAGCGGAGTGTATACTGGCTTACTATGTTGGCACTGA<br>TGAGGGTGTCAGTGAAGTGCTTCATGTGGCAGGAGAAAAA<br>AGGCTGCACCGGTGCGTCAGCAGAATATGTGATACAGGAT<br>ATATTCCGCTTCCTCGCTCACTGACTCGCTACGCTCGGTCG<br>TTCGACTGCGGCGAGCGGAAATGGCTTACGAACGGGGCG<br>GAGATTTCCTGGAAGATGCCAGGAAGATACTTAACAGGGAA<br>GTGAGAGGGCCGCGGCAAAGCCGTTTTTCCATAGGCTCCG<br>CCCCCCTGACAAGCATCACGAAATCTGACGCTCAAATCAGT<br>GGTGGCGAAACCCGACAGGACTATAAAGATACCAGGCGTT<br>TCCCCCTGGCGGCTCCCTCGTGCGCTCTCCTGTTCCTGCC<br>TTTCGGTTTACCGGTGTCATTCCGCTGTTATGGCCGCGTTT<br>GTCTCATTCCACGCCTGACACTCAGTTCCGGGTAGGCAGTT<br>CGCTCCAAGCTGGACTGTATGCACGAACCCCCCGTTCAGT<br>CCGACCGCTGCGCCTTATCCGGTAACTATCGTCTTGAGTCC<br>AACCCGGAAAGACATGCAAAAGCACCACTGGCAGCAGCCA<br>CTGGTAATTGATTTAGAGGAGTTAGTCTTGAAGTCATGCGC<br>CGGTTAAGGCTAAACTGAAAGGACAAGTTTTGGTGACTGCG<br>CTCCTCCAAGCCAGTTACCTCGGTTCAAAGAGTTGGTAGCT<br>CAGAGAACCTTCGAAAAACCGCCCTGCAAGGCGGTTTTTTC<br>GTTTTCAGAGCAAGAGATTACGCGCAGACCAAAACGATCTC<br>AAGAAGATCATCTTATTAATCAGATAAAATATTT |
| ColE1 origin of replication (BamHI-BglII scar) | GGCCGCGTTGCTGGCGTTTTTCCACAGGCTCCGCCCCCCT<br>GACGAGCATCACAAAAATCGACGCTCAAGTCAGAGGTGGC<br>GAAACCCGACAGGACTATAAAGATACCAGGCGTTTCCCCCT<br>GGAAGCTCCCTCGTGCGCTCTCCTGTTCCGACCCTGCCGC<br>TTACCGGATACCTGTCCGCCTTTCTCCCTTCGGGAAGCGTG |

| | |
|---|---|
| | GCGCTTTCTCATAGCTCACGCTGTAGGTATCTCAGTTCGGT<br>GTAGGTCGTTCGCTCCAAGCTGGGCTGTGTGCACGAACCC<br>CCCGTTCAGCCCGACCGCTGCGCCTTATCCGGTAACTATC<br>GTCTTGAGTCCAACCCGGTAAGACACGACTTATCGCCACTG<br>GCAGCAGCCACTGGTAACAGGATTAGCAGAGCGAGGTATG<br>TAGGCGGTGCTACAGAGTTCTTGAAGTGGTGGCCTAACTAC<br>GGCTACACTAGAAGAACAGTATTTGGTATCTGCGCTCTGCT<br>GAAGCCAGTTACCTTCGGAAAAAGAGTTGGTAGCTCTTGAT<br>CCGGCAAACAAACCACCGCTGGTAGCGGTGGTTTTTTTGTT<br>TGCAAGCAGCAGATTACGCGCAGAAAAAAAGGATCTCAAGA<br>AGATCCTTTGATCTTTTCTACGGGGTCTGACGCTCAGTGGA<br>ACGAAAACTCACGTTAAGGGATTTTGGTCATGA |
| CDF origin of replication | GCGCTGCGGACACATACAAAGTTACCCACAGATTCCGTGG<br>ATAAGCAGGGGACTAACATGTGAGGCAAAACAGCAGGGCC<br>GCGCCGGTGGCGTTTTTCCATAGGCTCCGCCCTCCTGCCA<br>GAGTTCACATAAACAGACGCTTTTCCGGTGCATCTGTGGGA<br>GCCGTGAGGCTCAACCATGAATCTGACAGTACGGGCGAAA<br>CCCGACAGGACTTAAAGATCCCCACCGTTTCCGGCGGGTC<br>GCTCCCTCTTGCGCTCTCCTGTTCCGACCCTGCCGTTTACC<br>GGATACCTGTTCCGCCTTTCTCCCTTACGGGAAGTGTGGCG<br>CTTTCTCATAGCTCACACACTGGTATCTCGGCTCGGTGTAG<br>GTCGTTCGCTCCAAGCTGGGCTGTAAGCAAGAACTCCCCG<br>TTCAGCCCGACTGCTGCGCCTTATCCGGTAACTGTTCACTT<br>GAGTCCAACCCGGAAAAGCACGGTAAAACGCCACTGGCAG<br>CAGCCATTGGTAACTGGGAGTTCGCAGAGGATTTGTTTAGC<br>TAAACACGCGGTTGCTCTTGAAGTGTGCGCCAAAGTCCGG<br>CTACACTGGAAGGACAGATTTGGTTGCTGTGCTCTGCGAAA<br>GCCAGTTACCACGGTTAAGCAGTTCCCCAACTGACTTAACC<br>TTCGATCAAACCACCTCCCCAGGTGGTTTTTTCGTTTACAG<br>GGCAAAAGATTACGCGCAGAAAAAAAGGATCTCAAGAAGAT<br>CCTTTGATC |

**Supplementary Table 2.** Summary of plasmids used in this study.

| Plasmid ID | Relevant Figure(s) | Plasmid organization |
|---|---|---|
| pJBL7201 | 1,2,3, S4, S6, S7 | J23101 – Target 1 – sfGFP – TrrnB – CmR – p15A |
| pJBL7202 | 1,2,3, S4 | J23101 – Target 2 – sfGFP – TrrnB – CmR – p15A |
| pJBL7203 | 1,2,3, S4 | J23101 – Target 3 – sfGFP – TrrnB – CmR – p15A |
| pJBL7204 | 1,2,3, S4 | J23101 – Target 4 – sfGFP – TrrnB – CmR – p15A |
| pJBL7205 | 1,2,3, S4, S6 | J23101 – Target 5 – sfGFP – TrrnB – CmR – p15A |
| pJBL7206 | 1,2,3, S4 | J23101 – Target 6 – sfGFP – TrrnB – CmR – p15A |
| pJBL7207 | 1,3, S4 | J23101 – Target 7 – sfGFP – TrrnB – CmR – p15A |
| pJBL7208 | 1,3, S4 | J23101 – Target 8 – sfGFP – TrrnB – CmR – p15A |

| | | |
|---|---|---|
| pJBL7209 | 1,3, S4 | J23101 – Target 9 – sfGFP – TrrnB – CmR – p15A |
| pJBL7210 | 1,3, S4 | J23101 – Target 10 – sfGFP – TrrnB – CmR – p15A |
| pJBL7211 | 1,3, S4 | J23101 – Target 11 – sfGFP – TrrnB – CmR – p15A |
| pJBL7212 | 1,3, S4 | J23101 – Target 12 – sfGFP – TrrnB – CmR – p15A |
| pJBL7213 | 1,3, S4, S6 | J23101 – Target 13 – sfGFP – TrrnB – CmR – p15A |
| pJBL7214 | 1,3, S4 | J23101 – Target 14 – sfGFP – TrrnB – CmR – p15A |
| pJBL7215 | 1,3, S4 | J23101 – Target 15 – sfGFP – TrrnB – CmR – p15A |
| pJBL7216 | 1,3, S4 | J23101 – Target 16 – sfGFP – TrrnB – CmR – p15A |
| pJBL7217 | 1,3, S4 | J23101 – Target 17 – sfGFP – TrrnB – CmR – p15A |
| pJBL7218 | 1,3, S4, S6 | J23101 – Target 18 – sfGFP – TrrnB – CmR – p15A |
| pJBL7219 | 1,3, S4 | J23101 – Target 19 – sfGFP – TrrnB – CmR – p15A |
| pJBL7220 | 1,3, S4 | J23101 – Target 20 – sfGFP – TrrnB – CmR – p15A |
| pJBL7221 | 1,3, S4 | J23101 – Target 21 – sfGFP – TrrnB – CmR – p15A |
| pJBL7222 | 1,3, S4 | J23101 – Target 22 – sfGFP – TrrnB – CmR – p15A |
| pJBL7223 | 1,3, S4 | J23101 – Target 23 – sfGFP – TrrnB – CmR – p15A |
| pJBL7224 | 1,2,3, S4, S5 | J23101 – Target 24 – sfGFP – TrrnB – CmR – p15A |
| pJBL7225 | 1,2,3, S4, S5 | J23101 – Target 25 – sfGFP – TrrnB – CmR – p15A |
| pJBL7226 | 1,2,3, S4, S5 | J23101 – Target 26 – sfGFP – TrrnB – CmR – p15A |
| pJBL7227 | 1,2,3,6, S4, S5 | J23101 – Target 27 – sfGFP – TrrnB – CmR – p15A |
| pJBL7228 | 1,2,3, S4, S5 | J23101 – Target 28 – sfGFP – TrrnB – CmR – p15A |
| pJBL7229 | 1,2,3, S4, S5 | J23101 – Target 29 – sfGFP – TrrnB – CmR – p15A |
| pJBL7230 | 1,2,3, S4, S5 | J23101 – Target 30 – sfGFP – TrrnB – CmR – p15A |
| pJBL7231 | 1,2,3, S4, S5 | J23101 – Target 31 – sfGFP – TrrnB – CmR – p15A |
| pJBL7232 | 1,2,3, S4, S5 | J23101 – Target 32 – sfGFP – TrrnB – CmR – p15A |
| pJBL7233 | 1,2,3, S4, S5 | J23101 – Target 33 – sfGFP – TrrnB – CmR – p15A |
| pJBL7234 | 1,2,3, S4, S5 | J23101 – Target 34 – sfGFP – TrrnB – CmR – p15A |
| pJBL7235 | 1,2,3, S4, S5 | J23101 – Target 35 – sfGFP – TrrnB – CmR – p15A |
| pJBL7236 | 1,2,3, S4, S5 | J23101 – Target 36 – sfGFP – TrrnB – CmR – p15A |
| pJBL7237 | 1,2,3, S4, S5 | J23101 – Target 37 – sfGFP – TrrnB – CmR – p15A |
| pJBL7238 | 1,2,3, S4, S5 | J23101 – Target 38 – sfGFP – TrrnB – CmR – p15A |
| pJBL7239 | 1,2,3, S4, S5 | J23101 – Target 39 – sfGFP – TrrnB – CmR – p15A |
| pJBL7240 | 1,2,3, S4, S5 | J23101 – Target 40 – sfGFP – TrrnB – CmR – p15A |
| pJBL7241 | 1,2,3, S4, S5 | J23101 – Target 41 – sfGFP – TrrnB – CmR – p15A |
| pJBL7242 | 1,3, S4 | J23101 – Target 42 – sfGFP – TrrnB – CmR – p15A |
| pJBL7243 | 1,3, S4 | J23101 – Target 43 – sfGFP – TrrnB – CmR – p15A |
| pJBL7244 | 1,3, S4 | J23101 – Target 44 – sfGFP – TrrnB – CmR – p15A |
| pJBL7245 | 1,3, S4 | J23101 – Target 45 – sfGFP – TrrnB – CmR – p15A |
| pJBL7246 | 1,3, S4 | J23101 – Target 46 – sfGFP – TrrnB – CmR – p15A |
| pJBL7247 | 1,2,3, S4, S5 | J23101 – Trigger 1 – T500 – ampR – ColE1 |
| pJBL7248 | 1,2, S4, S5 | J23101 – Trigger 2 – T500 – ampR – ColE1 |
| pJBL7249 | 1,2, S4, S5 | J23101 – Trigger 3 – T500 – ampR – ColE1 |

| | | |
|---|---|---|
| pJBL7250 | 1,2, S4, S5 | J23101 – Trigger 4 – T500 – ampR – ColE1 |
| pJBL7251 | 1,2, S4, S5 | J23101 – Trigger 5 – T500 – ampR – ColE1 |
| pJBL7252 | 1,2, S4, S5 | J23101 – Trigger 6 – T500 – ampR – ColE1 |
| pJBL7253 | 1, S4 | J23101 – Trigger 7 – T500 – ampR – ColE1 |
| pJBL7254 | 1, S4 | J23101 – Trigger 8 – T500 – ampR – ColE1 |
| pJBL7255 | 1, S4 | J23101 – Trigger 9 – T500 – ampR – ColE1 |
| pJBL7256 | 1, S4 | J23101 – Trigger 10 – T500 – ampR – ColE1 |
| pJBL7257 | 1, S4 | J23101 – Trigger 11 – T500 – ampR – ColE1 |
| pJBL7258 | 1, S4 | J23101 – Trigger 12 – T500 – ampR – ColE1 |
| pJBL7259 | 1, S4 | J23101 – Trigger 13 – T500 – ampR – ColE1 |
| pJBL7260 | 1, S4 | J23101 – Trigger 14 – T500 – ampR – ColE1 |
| pJBL7261 | 1, S4 | J23101 – Trigger 15 – T500 – ampR – ColE1 |
| pJBL7262 | 1, S4 | J23101 – Trigger 16 – T500 – ampR – ColE1 |
| pJBL7263 | 1, S4 | J23101 – Trigger 17 – T500 – ampR – ColE1 |
| pJBL7264 | 1, S4 | J23101 – Trigger 18 – T500 – ampR – ColE1 |
| pJBL7265 | 1, S4 | J23101 – Trigger 19 – T500 – ampR – ColE1 |
| pJBL7266 | 1, S4 | J23101 – Trigger 20 – T500 – ampR – ColE1 |
| pJBL7267 | 1, S4 | J23101 – Trigger 21 – T500 – ampR – ColE1 |
| pJBL7268 | 1, S4 | J23101 – Trigger 22 – T500 – ampR – ColE1 |
| pJBL7269 | 1, S4 | J23101 – Trigger 23 – T500 – ampR – ColE1 |
| pJBL7270 | 1,2, S4, S5 | J23101 – LASO 24 – T500 – ampR – ColE1 |
| pJBL7271 | 1,2, S4, S5 | J23101 – LASO 25 – T500 – ampR – ColE1 |
| pJBL7272 | 1,2, S4, S5 | J23101 – LASO 26 – T500 – ampR – ColE1 |
| pJBL7273 | 1,2, S4, S5 | J23101 – LASO 27 – T500 – ampR – ColE1 |
| pJBL7274 | 1,2, S4, S5 | J23101 – LASO 28 – T500 – ampR – ColE1 |
| pJBL7275 | 1,2, S4, S5 | J23101 – LASO 29 – T500 – ampR – ColE1 |
| pJBL7276 | 1,2, S4, S5 | J23101 – LASO 30 – T500 – ampR – ColE1 |
| pJBL7277 | 1,2, S4, S5 | J23101 – LASO 31 – T500 – ampR – ColE1 |
| pJBL7278 | 1,2, S4, S5 | J23101 – LASO 32 – T500 – ampR – ColE1 |
| pJBL7279 | 1,2, S4, S5 | J23101 – LASO 33 – T500 – ampR – ColE1 |
| pJBL7280 | 1,2, S4, S5 | J23101 – LASO 34 – T500 – ampR – ColE1 |
| pJBL7281 | 1,2, S4, S5 | J23101 – LASO 35 – T500 – ampR – ColE1 |
| pJBL7282 | 1,2, S4, S5 | J23101 – LASO 36 – T500 – ampR – ColE1 |
| pJBL7283 | 1,2, S4, S5 | J23101 – LASO 37 – T500 – ampR – ColE1 |
| pJBL7284 | 1,2, S4, S5 | J23101 – LASO 38 – T500 – ampR – ColE1 |
| pJBL7285 | 1,2, S4, S5 | J23101 – LASO 39 – T500 – ampR – ColE1 |
| pJBL7286 | 1,2, S4, S5 | J23101 – LASO 40 – T500 – ampR – ColE1 |
| pJBL7287 | 1,2, S4, S5 | J23101 – LASO 41 – T500 – ampR – ColE1 |
| pJBL7288 | 1, S4 | J23101 – LASO 42 – T500 – ampR – ColE1 |
| pJBL7289 | 1, S4 | J23101 – LASO 43 – T500 – ampR – ColE1 |
| pJBL7290 | 1, S4 | J23101 – LASO 44 – T500 – ampR – ColE1 |

| | | |
|---|---|---|
| pJBL7291 | 1, S4 | J23101 – LASO 45 – T500 – ampR – ColE1 |
| pJBL7292 | 1, S4 | J23101 – LASO 46 – T500 – ampR – ColE1 |
| pJBL7293 | 3, S6 | J23101 – Target 1 (+ 4nt) – sfGFP – TrrnB – CmR – p15A |
| pJBL7294 | 3,7, S6 | J23101 – Target 1 (+ 8nt) – sfGFP – TrrnB – CmR – p15A |
| pJBL7295 | 3, S6 | J23101 – Target 5 (+ 4nt) – sfGFP – TrrnB – CmR – p15A |
| pJBL7296 | 3, S6 | J23101 – Target 5 (+ 8nt) – sfGFP – TrrnB – CmR – p15A |
| pJBL7297 | 3, S6 | J23101 – Target 13 (+ 4nt) – sfGFP – TrrnB – CmR – p15A |
| pJBL7298 | 3, S6 | J23101 – Target 13 (+ 8nt) – sfGFP – TrrnB – CmR – p15A |
| pJBL7299 | 3, S6 | J23101 – Target 18 (+ 4nt) – sfGFP – TrrnB – CmR – p15A |
| pJBL7300 | 3, S6 | J23101 – Target 18 (+ 8nt) – sfGFP – TrrnB – CmR – p15A |
| pJBL7301 | 3,7, S7 | J23119 – Trigger 1 – T500 – ampR – ColE1 |
| pJBL7302 | 4(a,d), S8, S9 | J23119(SpeI) – asRNA (targeting mRFP) – T500 – ampR – ColE1 |
| pJBL7303 | 4(b,e,f,h), S8, S9 | J23119(SpeI) – LASO (targeting mRFP, variant 1) – T500 – ampR – ColE1 |
| pJBL7304 | 4(f,h), S8, S9 | J23119(SpeI) – LASO (targeting mRFP, variant 2) – T500 – ampR – ColE1 |
| pJBL7305 | 4(f,h), S8, S9 | J23119(SpeI) – LASO (targeting mRFP, variant 3) – T500 – ampR – ColE1 |
| pJBL7306 | 4(f,h), S8, S9 | J23119(SpeI) – LASO (targeting mRFP, variant 4) – T500 – ampR – ColE1 |
| pJBL7307 | 4(f,h), S8, S9 | J23119(SpeI) – LASO (targeting mRFP, variant 5) – T500 – ampR – ColE1 |
| pJBL7308 | 4(f,h), S8, S9 | J23119(SpeI) – LASO (targeting mRFP, variant 6) – T500 – ampR – ColE1 |
| pJBL7309 | 4(f,h), S8, S9 | J23119(SpeI) – LASO (targeting mRFP, variant 7) – T500 – ampR – ColE1 |
| pJBL7310 | 4(f,h), S8, S9 | J23119(SpeI) – LASO (targeting mRFP, variant 8) – T500 – ampR – ColE1 |
| pJBL7311 | 4(f,h), S8, S9 | J23119(SpeI) – LASO (targeting mRFP, variant 9) – T500 – ampR – ColE1 |
| pJBL7312 | 4(f,h), S8, S9 | J23119(SpeI) – LASO (targeting mRFP, variant 10) – T500 – ampR – ColE1 |
| pJBL7313 | 4(f,h), S8, S9 | J23119(SpeI) – LASO (targeting mRFP, variant 11) – T500 – ampR – ColE1 |

| | | |
|---|---|---|
| pJBL7314 | 4(f,h), S8, S9 | J23119(SpeI) – LASO (targeting mRFP, variant 12) – T500 – ampR – ColE1 |
| pJBL7315 | 4(f,h), S8, S9 | J23119(SpeI) – LASO (targeting mRFP, variant 13) – T500 – ampR – ColE1 |
| pJBL7316 | 4(f,h), S8, S9 | J23119(SpeI) – LASO (targeting mRFP, variant 14) – T500 – ampR – ColE1 |
| pJBL7317 | 4(f,h), S8, S9 | J23119(SpeI) – LASO (targeting mRFP, variant 15) – T500 – ampR – ColE1 |
| pJBL7318 | 4(f,h), S8, S9 | J23119(SpeI) – LASO (targeting mRFP, variant 16) – T500 – ampR – ColE1 |
| pJBL7319 | 4(f,h), S8, S9 | J23119(SpeI) – LASO (targeting mRFP, variant 17) – T500 – ampR – ColE1 |
| pJBL7320 | 4(f,h), S8, S9 | J23119(SpeI) – LASO (targeting mRFP, variant 18) – T500 – ampR – ColE1 |
| pJBL7321 | 4(f,h), S8, S9 | J23119(SpeI) – LASO (targeting mRFP, variant 19) – T500 – ampR – ColE1 |
| pJBL7322 | 4(f,h), S8, S9 | J23119(SpeI) – LASO (targeting mRFP, variant 20) – T500 – ampR – ColE1 |
| pJBL7323 | 4(f,h), S8, S9 | J23119(SpeI) – LASO (targeting mRFP, variant 21) – T500 – ampR – ColE1 |
| pJBL7324 | 4(f,h), S8, S9 | J23119(SpeI) – LASO (targeting mRFP, variant 22) – T500 – ampR – ColE1 |
| pJBL7325 | 4(f,h), S8, S9 | J23119(SpeI) – LASO (targeting mRFP, variant 23) – T500 – ampR – ColE1 |
| pJBL7326 | 4(f,h), S8, S9 | J23119(SpeI) – LASO (targeting mRFP, variant 24) – T500 – ampR – ColE1 |
| pJBL7327 | 4(f,h), S8, S9 | J23119(SpeI) – LASO (targeting mRFP, variant 25) – T500 – ampR – ColE1 |
| pJBL7328 | 4(f,h), S8, S9 | J23119(SpeI) – LASO (targeting mRFP, variant 26) – T500 – ampR – ColE1 |
| pJBL7329 | 4(f,h), S8, S9 | J23119(SpeI) – LASO (targeting mRFP, variant 27) – T500 – ampR – ColE1 |
| pJBL7330 | 4(f,h), S8, S9 | J23119(SpeI) – LASO (targeting mRFP, variant 28) – T500 – ampR – ColE1 |
| pJBL7331 | 4(f,h), S8, S9 | J23119(SpeI) – LASO (targeting mRFP, variant 29) – T500 – ampR – ColE1 |
| pJBL7332 | 4(f,h), S8, S9 | J23119(SpeI) – LASO (targeting mRFP, variant 30) – T500 – ampR – ColE1 |
| pJBL7333 | 4(f,h), S8, S9 | J23119(SpeI) – LASO (targeting mRFP, variant 31) – T500 – ampR – ColE1 |
| pJBL7334 | 4(f,h), S8, S9 | J23119(SpeI) – LASO (targeting mRFP, variant 32) – T500 – ampR – ColE1 |
| pJBL7335 | 4(f,h), S8, S9 | J23119(SpeI) – LASO (targeting mRFP, variant 33) – T500 – ampR – ColE1 |

| | | |
|---|---|---|
| pJBL7336 | 4(f,h), S8, S9 | J23119(SpeI) – LASO (targeting mRFP, variant 34) – T500 – ampR – ColE1 |
| pJBL7337 | 4(f,h), S8, S9 | J23119(SpeI) – LASO (targeting mRFP, variant 35) – T500 – ampR – ColE1 |
| pJBL7338 | 4(f,h), S8, S9 | J23119(SpeI) – LASO (targeting mRFP, variant 36) – T500 – ampR – ColE1 |
| pJBL7339 | 4(f,h), S8, S9 | J23119(SpeI) – LASO (targeting mRFP, variant 37) – T500 – ampR – ColE1 |
| pJBL7340 | 4(f,h), S8, S9 | J23119(SpeI) – LASO (targeting mRFP, variant 38) – T500 – ampR – ColE1 |
| pJBL7341 | 4(f,h), S8, S9 | J23119(SpeI) – LASO (targeting mRFP, variant 39) – T500 – ampR – ColE1 |
| pJBL7342 | 4(f,h), S8, S9 | J23119(SpeI) – LASO (targeting mRFP, variant 40) – T500 – ampR – ColE1 |
| pJBL7343 | 4(f,h), S8, S9 | J23119(SpeI) – LASO (targeting mRFP, variant 41) – T500 – ampR – ColE1 |
| pJBL7344 | 4(f,h), S8, S9 | J23119(SpeI) – LASO (targeting mRFP, variant 42) – T500 – ampR – ColE1 |
| pJBL7345 | 4(f,h), S8, S9 | J23119(SpeI) – LASO (targeting mRFP, variant 43) – T500 – ampR – ColE1 |
| pJBL7346 | 4(f,h), S8, S9 | J23119(SpeI) – LASO (targeting mRFP, variant 44) – T500 – ampR – ColE1 |
| pJBL7347 | 4(f,h), S8, S9 | J23119(SpeI) – LASO (targeting mRFP, variant 45) – T500 – ampR – ColE1 |
| pJBL7348 | 4(f,h), S8, S9 | J23119(SpeI) – LASO (targeting mRFP, variant 46) – T500 – ampR – ColE1 |
| pJBL7349 | 4(f,h), S8, S9 | J23119(SpeI) – LASO (targeting mRFP, variant 47) – T500 – ampR – ColE1 |
| pJBL7350 | 4(f,h), S8, S9 | J23119(SpeI) – LASO (targeting mRFP, variant 48) – T500 – ampR – ColE1 |
| pJBL7351 | 4(f,h), S8, S9 | J23119(SpeI) – LASO (targeting mRFP, variant 49) – T500 – ampR – ColE1 |
| pJBL7352 | 4(f,h), S8, S9 | J23119(SpeI) – LASO (targeting mRFP, variant 50) – T500 – ampR – ColE1 |
| pJBL7353 | 4(f,h), S8, S9 | J23119(SpeI) – LASO (targeting mRFP, variant 51) – T500 – ampR – ColE1 |
| pJBL7354 | 4(f,h), S8, S9 | J23119(SpeI) – LASO (targeting mRFP, variant 52) – T500 – ampR – ColE1 |
| pJBL6654 | 5(c,d), S10 | J23101 – *gnd* (126nt) – sfGFP – TrrnB – CmR – p15A |
| pJBL6655 | 5(c,d), S10 | J23101 – *gmd* (151nt) – sfGFP – TrrnB – CmR – p15A |
| pJBL6662 | 5(c,d), S10 | J23101 – *murC* (151nt) – sfGFP – TrrnB – CmR – p15A |

| | | |
|---|---|---|
| pJBL6663 | 5(c,d), S10 | J23101 – *pfkA* (101nt) – sfGFP – TrrnB – CmR – p15A |
| pJBL6664 | 5(c,d), S10 | J23101 – *zwf* (151nt) – sfGFP – TrrnB – CmR – p15A |
| pJBL6628 | 5(c,d), S10 | J23101 – LASO (targeting *gnd*) – T500 – ampR – ColE1 |
| pJBL6614 | 5(c,d), S10 | J23101 – LASO (targeting *gmd*) – T500 – ampR – ColE1 |
| pJBL6633 | 5(c,d), S10 | J23101 – LASO (targeting *murC*) – T500 – ampR – ColE1 |
| pJBL6634 | 5(c,d), S10 | J23101 – LASO (targeting *pfkA*) – T500 – ampR – ColE1 |
| pJBL6635 | 5(c,d), S10 | J23101 – LASO (targeting *zwf*) – T500 – ampR – ColE1 |
| pJBL7355 | 6, S11 | J23101 – Target 27 (RBS = TAAGGCTC, ATG start codon) – sfGFP – TrrnB – CmR – p15A |
| pJBL7356 | 6, S11 | J23101 – Target 27 (RBS = TGAAGGGG, ATG start codon) – sfGFP – TrrnB – CmR – p15A |
| pJBL7357 | 6, S11 | J23101 – Target 27 (RBS = AAGGACCC, ATG start codon) – sfGFP – TrrnB – CmR – p15A |
| pJBL7358 | 6, S11 | J23101 – Target 27 (RBS = TGAAGGGG, ATG start codon) – sfGFP – TrrnB – CmR – p15A |
| pJBL7359 | 6, S11 | J23101 – Target 27 (RBS = GAGGCGA, ATG start codon) – sfGFP – TrrnB – CmR – p15A |
| pJBL7360 | 6, S11 | J23101 – Target 27 (RBS = GGTGGGCA, ATG start codon) – sfGFP – TrrnB – CmR – p15A |
| pJBL7361 | 6, S11 | J23101 – Target 27 (RBS = AGAAGAGA, ATG start codon) – sfGFP – TrrnB – CmR – p15A |
| pJBL7362 | 6, S11 | J23101 – Target 27 (RBS = AAGAGAGG, ATG start codon) – sfGFP – TrrnB – CmR – p15A |
| pJBL7363 | 6, S11 | J23101 – Target 27 (RBS = AAGTGAAA, ATG start codon) – sfGFP – TrrnB – CmR – p15A |
| pJBL7364 | 6, S11 | J23101 – Target 27 (RBS = AGGCCGAC, ATG start codon) – sfGFP – TrrnB – CmR – p15A |
| pJBL7365 | 6, S11 | J23101 – Target 27 (RBS = AGAGGATG, ATG start codon) – sfGFP – TrrnB – CmR – p15A |
| pJBL7366 | 6, S11 | J23101 – Target 27 (RBS = ACACGAAC, ATG start codon) – sfGFP – TrrnB – CmR – p15A |
| pJBL7367 | 6, S11 | J23101 – Target 27 (RBS = CAGGT, ATG start codon) – sfGFP – TrrnB – CmR – p15A |
| pJBL7368 | 6, S11 | J23101 – Target 27 (RBS = AGGCAG, ATG start codon) – sfGFP – TrrnB – CmR – p15A |
| pJBL7369 | 6, S11 | J23101 – Target 27 (RBS = CACCCCCC, ATG start codon) – sfGFP – TrrnB – CmR – p15A |

| | | |
|---|---|---|
| pJBL7370 | 6, S11 | J23101 – Target 27 (RBS = GCGTGGGA, GTG start codon) – sfGFP – TrrnB – CmR – p15A |
| pJBL7371 | 6, S11 | J23101 – Target 27 (RBS = TCTTGAGA, ATG start codon) – sfGFP – TrrnB – CmR – p15A |
| pJBL7372 | 6, S11 | J23101 – Target 27 (RBS = GTGAGAAG, ATG start codon) – sfGFP – TrrnB – CmR – p15A |
| pJBL7373 | 6, S11 | J23101 – Target 27 (RBS = AAAGGCAA, ATG start codon) – sfGFP – TrrnB – CmR – p15A |
| pJBL7374 | 6, S11 | J23101 – Target 27 (RBS = GGGTGGTA, ATG start codon) – sfGFP – TrrnB – CmR – p15A |
| pJBL7375 | 6, S11 | J23101 – Target 27 (RBS = TCAATAGG, ATG start codon) – sfGFP – TrrnB – CmR – p15A |
| pJBL7376 | 6, S11 | J23101 – Target 27 (RBS = ACCCTTAT, ATG start codon) – sfGFP – TrrnB – CmR – p15A |
| pJBL7377 | 6, S11 | J23101 – Target 27 (RBS = ATGGG, GTG start codon) – sfGFP – TrrnB – CmR – p15A |
| pJBL7378 | 6, S11 | J23101 – Target 27 (RBS = CAACTAAC, GTG start codon) – sfGFP – TrrnB – CmR – p15A |
| pJBL7379 | 6, S11 | J23101 – Target 27 (RBS = TACCATTA, ATG start codon) – sfGFP – TrrnB – CmR – p15A |
| pJBL7380 | 6, S11 | J23101 – Target 27 (RBS = GAGACAG, ATG start codon) – sfGFP – TrrnB – CmR – p15A |
| pJBL7381 | 6, S11 | J23101 – Target 27 (RBS = GGCATTGG, ATG start codon) – sfGFP – TrrnB – CmR – p15A |
| pJBL7382 | 6, S11 | J23101 – Target 27 (RBS = GAAGGGAC, ATG start codon) – sfGFP – TrrnB – CmR – p15A |
| pJBL7383 | 6, S11 | J23101 – Target 27 (RBS = CGGCCCAG, ATG start codon) – sfGFP – TrrnB – CmR – p15A |
| pJBL7384 | 6, S11 | J23101 – Target 27 (RBS = TAAAGGAC, GTG start codon) – sfGFP – TrrnB – CmR – p15A |
| pJBL7385 | 6, S11 | J23119(SpeI) – LASO 27 – T500 – ampR – ColE1 |
| pJBL7386 | 7, S12 | J23150 – Toehold switch 1 – mRFP – Double terminator – specR - CDF |
| pJBL7387 | S7 | J23150 – Target 1 – sfGFP – TrrnB – CmR – p15A |
| pJBL7388 | S7 | J23101 – Target 1 – GFPmut3bASV – TrrnB – CmR – p15A |

**Supplementary Table 3.** Sequences of toehold repressor target designs used in this study.

| Description | Sequence |
|---|---|
| Target 1 | AGAAAGAAGAAGAGATGCGGAAGAGAGAAATATAACACAACAATACGTATATTTATATCTTCCGCAAAGAAGAGGAGAGGAAGAATGAAATATACGAGACAACAAACAATAAACAGC |
| Target 2 | TAGATGAAACGAGAACAGCGCGAAGTAATTGAACCTAATCGATATCATTTCAATGACTTCGCGCTGGAACAGAGGAGACGCGAAATGATTGAAATGAACCTGGCGGCAGCGCAAAAG |
| Target 3 | AGAGAAAGAAGAAGAACGCGAAAGAAGAAGATACAAACAACACAATCTTATCTTATGCTTTCGCGTAAAGAGAGGAGACGAAAGATGAAGATAAGAACAAACAAACAAACAAACAAC |
| Target 4 | ATCACTTATTGTCGTTGCTGTATGTCTGTAAATCACTTCTAAACATCGATTTACTTCCATACAGCATAAGAGAGGAGATGTATGATGGTAAATCGACACAAAGCTATAAATCAAATC |
| Target 5 | AATAACGAGTGACAAGGGTGATGAGTGAAGCAGAATGATTAAGATAGGCTGCTTAGATCATCACCCGGATAGAGGAGATGATGAATGAAGCAGCCTAACCTGGCGGCAGCGCAAAAG |
| Target 6 | TAGAATTTGATACTTTCCCGACGCTTAAGAAATATATAAATCCTCCTTATTTCTGTCGCGTCGGGACACAAGAGGAGACGACGCATGAGAAATAAGAACAACAAACAAACAACAAAC |
| Target 7 | GAGAGTGAAAGTGAACGCGAGGGTGAATAGAATAGATGAGGATATACCTCATCTATTCATTTCACCCTCGCGAAATAGAGGAGACGCGAGATGGAAATGAATAGAAACCTGGCGGCAGCGCAAAAG |
| Target 8 | GATTGTAACGTAAGTCCGAGTGTAGTGTAAGTAAGCTGAGGACAATCCTCAGCTTACTATCACTACACTCGGAAATAGAGGAGACCGAGTATGGTGATAGTAAGCAACCTGGCGGCAGCGCAAAAG |
| Target 9 | AGTAAATGTAGTAAACGCGAGCGAATAAGTAAGAATGAGGTAGACGACCTCATTCTTAGTTATTCGCTCGCGAAATAGAGGAGACGCGAGATGATAACTAAGAATAACCTGGCGGCAGCGCAAAAG |
| Target 10 | GGGAATGAAGTGAACGGCTGAAGTAGAATAAGGACAACTCAACTATACCCTTATGTGACTTCAGCCAAGTAGAGGAGATGAAGTATGATAAGGGTAAACCTGGCGGCAGCGCAAAAG |
| Target 11 | GAGACATATACAAAGCAGCAAAGTTAAGTACTAGAGATAAACGACATGTAGTACGTTACTTTGCTGGGACAGAGGAGACAAAGTATGGTACTACATAACCTGGCGGCAGCGCAAAAG |
| Target 12 | GAATGGTAGAATAGTCGCGAAGGTAAACATGAGAATAGGCTTCAGAAGCCTATTCTCAGGTTTACCTTCGCGAGATAGAGGAGACGCGAAATGAAACCTGAGAATAACCTGGCGGCAGCGCAAAAG |

| | |
|---|---|
| Target 13 | GGGATGTAATGAGAACGGTAGGGTAGTAAAGATAGAGTGAGGCATACTCACTCTATCTCCACTACCCTACCGAAATAGAGGAGACGGTAGATGAGTGGAGATAGAAACCTGGCGGCAGCGCAAAAG |
| Target 14 | TATTACACATTCAAGCCCATTGCTAACGATGACACTTATGCCGTTTGCATAAGTGTCAAGGTTAGCAATGGGCAACAGAGGAGACCCATTATGAACCTTGACACTAACCTGGCGGCAGCGCAAAAG |
| Target 15 | ACAAAGATTGGTCGTACGATTACCGTTAGAATACACACACACACATCTTATTCTTTAGGTAATCGTGCAAAGAGGAGAATTACCATGAGAATAAGACAAACAAACAAACTAAACAAC |
| Target 16 | GGAGTGAAATGAAGACGGTAAGGTGGAGTGAATAGAGGTGAACTCATCACCTCTATTCTGTCCACCTTACCGAAACAGAGGAGACGGTAAATGGGACAGAATAGAAACCTGGCGGCAGCGCAAAAG |
| Target 17 | GGAGCATAAAGGTAAACGTAGTAATAGAGGCGATACGGGCATGAGATCTCGCCTGAATTACTACGTACATAGAGGAGATAGTAAATGAGGCGAGATAGAGACAGTAGGAGTGGAAGG |
| Target 18 | AGAACAAGTAAGATACGCCAAATTCAGCAACATAAGTAGGTACAGAACCTACTTATGTATCTGAATTTGGCGAAATAGAGGAGACGCCAAATGCAGATACATAAGAACCTGGCGGCAGCGCAAAAG |
| Target 19 | GGAGTAAGTAATGAACGCGAGGGTAAGTGAAATAGGTGAGGATCAACCTCACCTATTTGTCTTACCCTCGCGAAACAGAGGAGACGCGAGATGAAGACAAATAGGAACCTGGCGGCAGCGCAAAAG |
| Target 20 | GGTGAACAATAAGAACGGTAAGGTGAGGTAATGAATATGTAGATGATACATATTCATTTGCTCACCTTACCGGAATAGAGGAGACGGTAAATGGAGCAAATGAATAACCTGGCGGCAGCGCAAAAG |
| Target 21 | GAGATGTAATGAAGGCTGTGATGAGGTGAGACTAACGAAACATAACTCAGTCTCAAGTCATCACAGCAATAGAGGAGATGATGAATGGAGACTGAGAACCTGGCGGCAGCGCAAAAG |
| Target 22 | AGATGGAATAGAAATCGGAATGAGAGCAAGAATAAGTTAGATGACCTCTAACTTATTCATGCTCTCATTCCGACACAGAGGAGACGGAATATGAGCATGAATAAGAACCTGGCGGCAGCGCAAAAG |
| Target 23 | TGGCCCATAAGTAGTCGGTAGCGTGCTGTAGATGATGAATGATCCGCATTCATCATCTTCAGCACGCTACCGAAATAGAGGAGACGGTAGATGGCTGAAGATGATAACCTGGCGGCAGCGCAAAAG |
| Target 1, +4nt | AGAAAGAAGAAGAGATGCGGAAGAGAGAAATATAACACAACAATACGTATATTTATATCTTCCGCAAAGACTAGAGAGGAGAGGAAGAATGAAATATACGAGACAACAAACAATAAACAGC |

| | |
|---|---|
| Target 1, +8nt | AGAAAGAAGAAGAGATGCGGAAGAGAGAAATATAACACAACAATACGTATATTTATATCTTCCGCAAAGAACACTAATAGAGGAGAGGAAGAATGAAATATACGAGACAACAAACAATAAACAGC |
| Target 5, +4nt | AATAACGAGTGACAAGGGTGATGAGTGAAGCAGAATGATTAAGATAGGCTGCTTAGATCATCACCCGGATGATGAGAGGAGATGATGAATGAAGCAGCCTAACCTGGCGGCAGCGCAAAAG |
| Target 5, +8nt | AATAACGAGTGACAAGGGTGATGAGTGAAGCAGAATGATTAAGATAGGCTGCTTAGATCATCACCCGGATTTTAACTAAGAGGAGATGATGAATGAAGCAGCCTAACCTGGCGGCAGCGCAAAAG |
| Target 13, +4nt | GGGATGTAATGAGAACGGTAGGGTAGTAAAGATAGAGTGAGGCATACTCACTCTATCTCCACTACCCTACCGAAATTCTAAGAGGAGACGGTAGATGAGTGGAGATAGAAACCTGGCGGCAGCGCAAAAG |
| Target 13, +8nt | GGGATGTAATGAGAACGGTAGGGTAGTAAAGATAGAGTGAGGCATACTCACTCTATCTCCACTACCCTACCGAAATGGGCTATAAGAGGAGACGGTAGATGAGTGGAGATAGAAACCTGGCGGCAGCGCAAAAG |
| Target 18, +4nt | AGAACAAGTAAGATACGCCAAATTCAGCAACATAAGTAGGTACAGAACCTACTTATGTATCTGAATTTGGCGAAATGGTAAGAGGAGACGCCAAATGCAGATACATAAGAACCTGGCGGCAGCGCAAAAG |
| Target 18, +8nt | AGAACAAGTAAGATACGCCAAATTCAGCAACATAAGTAGGTACAGAACCTACTTATGTATCTGAATTTGGCGAAATATTATACAAGAGGAGACGCCAAATGCAGATACATAAGAACCTGGCGGCAGCGCAAAAG |

**Supplementary Table 4.** Sequences of toehold repressor trigger designs used in this study.

| Description | Sequence |
|---|---|
| Trigger 1 | ATATTTTTTTCTTCCGCATCTCTTCTTCTTTCT |
| Trigger 2 | TTCAATTACTTCGCGCTGTTCTCGTTTCATCTA |
| Trigger 3 | TGTCTTCTTCTTTCGCGTTCTTCTTCTTTCTCT |
| Trigger 4 | ATTTACGGACATACAGCAACGACAATAAGTGAT |
| Trigger 5 | CTGCTTCACTCATCACCCTTGTCACTCGTTATT |
| Trigger 6 | ATTTCTTAAGCGTCGGGAAAGTATCAAATTCTA |
| Trigger 7 | CCTCATCTATTCTATTCACCCTCGCGTTCACTTTCACTCTC |
| Trigger 8 | CCTCAGCTTACTTACACTACACTCGGACTTACGTTACAATC |
| Trigger 9 | ACCTCATTCTTACTTATTCGCTCGCGTTTACTACATTTACT |
| Trigger 10 | CCTTATTCTACTTCAGCCGTTCACTTCATTCCC |
| Trigger 11 | TAGTACTTAACTTTGCTGCTTTGTATATGTCTC |

| | |
|---|---|
| Trigger 12 | AGCCTATTCTCATGTTTACCTTCGCGACTATTCTACCATTC |
| Trigger 13 | CTCACTCTATCTTTACTACCCTACCGTTCTCATTACATCCC |
| Trigger 14 | GCATAAGTGTCATCGTTAGCAATGGGCTTGAATGTGTAATA |
| Trigger 15 | TATTCTAACGGTAATCGTACGACCAATCTTTGT |
| Trigger 16 | TCACCTCTATTCACTCCACCTTACCGTCTTCATTTCACTCC |
| Trigger 17 | TCGCCTCTATTACTACGTTTACCTTTATGCTCC |
| Trigger 18 | ACCTACTTATGTTGCTGAATTTGGCGTATCTTACTTGTTCT |
| Trigger 19 | CCTCACCTATTTCACTTACCCTCGCGTTCATTACTTACTCC |
| Trigger 20 | TACATATTCATTACCTCACCTTACCGTTCTTATTGTTCACC |
| Trigger 21 | AGTCTCACCTCATCACAGCCTTCATTACATCTC |
| Trigger 22 | TCTAACTTATTCTTGCTCTCATTCCGATTTCTATTCCATCT |
| Trigger 23 | CATTCATCATCTACAGCACGCTACCGACTACTTATGGGCCA |

**Supplementary Table 5.** Sequences of LASO target designs used in this study.

| Description | Sequence |
|---|---|
| Target 24 | AAATGATGGAATAAGAGAAGACAGAGGAGATAACATATGAT ACACAGCAACCTGGCGGCAGCGCAAAAG |
| Target 25 | ACGAATTTGGAAGTAAAGAAACAGAGGAGACATAACATGCT ACTACTCAACCTGGCGGCAGCGCAAAAG |
| Target 26 | ATAAGTTTGAATATGAAGAAACAGAGGAGATCAAAGATGTC ATTTCGCAACCTGGCGGCAGCGCAAAAG |
| Target 27 | AGGATCTAAATGTATACAGAACAGAGGAGACAATACATGAA CGAACGAAACCTGGCGGCAGCGCAAAAG |
| Target 28 | CAAATGATTAAGTGACAAGAACAGAGGAGATATAGAATGAA ACGACGAAACCTGGCGGCAGCGCAAAAG |
| Target 29 | CAATGTAATCAAACTCATAAACAGAGGAGATATCACATGAC TAAACGAAACCTGGCGGCAGCGCAAAAG |
| Target 30 | AGTTAAAGATGAGAAAGGAAACAGAGGAGATCACATATGTC ACAAGCGAACCTGGCGGCAGCGCAAAAG |
| Target 31 | AAGATAGATGATTGTGGACAACAGAGGAGACAGAATATGAA ACTAAGCAACCTGGCGGCAGCGCAAAAG |
| Target 32 | CGAATAGAAATGAAGCAGAAACAGAGGAGATATGACATGG ACACTAGCAACCTGGCGGCAGCGCAAAAG |
| Target 33 | AAACGTTAATCTCTGAACGAACAGAGGAGATACAGAATGAA AGCAAGCAACCTGGCGGCAGCGCAAAAG |
| Target 34 | TTCATTCATTCCATTCAGAAACAGAGGAGATATACAATGGA CAAGCAGAACCTGGCGGCAGCGCAAAAG |
| Target 35 | AGAGTAAGTAGAGGTGTAGAACAGAGGAGATATGCAATGG TAAGAGTGAACCTGGCGGCAGCGCAAAAG |
| Target 36 | ATAGATAAGATAAGACAGAAACAGAGGAGATATGCAATGAT AAACGAGAACCTGGCGGCAGCGCAAAAG |

| | |
|---|---|
| Target 37 | AAGATAGAAATTGATAAGAGACAGAGGAGACGGCAAATGAA ACTAACGAACCTGGCGGCAGCGCAAAAG |
| Target 38 | CGAATCACTTATTGTCGCGAACAGAGGAGATACAGAATGAC AAAGACGAACCTGGCGGCAGCGCAAAAG |
| Target 39 | CTTAATCTTACCTTCCAGAAACAGAGGAGATATCGGATGAG AACAAGCAACCTGGCGGCAGCGCAAAAG |
| Target 40 | CATCGAAAGTGTATGAATAAACAGAGGAGATAACTAATGCA TATCAGCAACCTGGCGGCAGCGCAAAAG |
| Target 41 | AGGACAAAGATTGGTAAAGAACAGAGGAGATCTAACATGAA TGAAACGAACCTGGCGGCAGCGCAAAAG |
| Target 42 | AAATATGAATTGATGAGAAGACAGAGGAGATCAAGAATGAA CAATCGAAACCTGGCGGCAGCGCAAAAG |
| Target 43 | TAAGTAGTAATAGATCAGAAACAGAGGAGATATAGAATGAG ACAATGGAACCTGGCGGCAGCGCAAAAG |
| Target 44 | TGTCAAAGTAGTAAGACAGAACAGAGGAGATATCGTATGAG AATCAGGAACCTGGCGGCAGCGCAAAAG |
| Target 45 | AAATGAATGAATTGTAAGAAACAGAGGAGACATAACATGAA CAAGCCTAACCTGGCGGCAGCGCAAAAG |
| Target 46 | TATGTAATTGATTTGCAAGAACAGAGGAGATATGAAATGAA CAGAAGCAACCTGGCGGCAGCGCAAAAG |

**Supplementary Table 6.** Sequences of LASO designs used in this study.

| Description | Sequence |
|---|---|
| LASO 24 | GCTGTGTATATGATGTTACTTATTCCATCATTT |
| LASO 25 | GAGTAGTAGATAGTTATGTACTTCCAAATTCGT |
| LASO 26 | GCGAAATGAATGCTTTGACATATTCAAACTTAT |
| LASO 27 | TCGTTCGTTATGGTATTGATACATTTAGATCCT |
| LASO 28 | TCGTCGTTTCTATCTATATCACTTAATCATTTG |
| LASO 29 | TCGTTTAGTTTAGTGATAAGTTTGATTACATTG |
| LASO 30 | CGCTTGTGAATGATGTGATTCTCATCTTTAGCT |
| LASO 31 | GCTTAGTTTATGATTCTGACAATCATCTATCTT |
| LASO 32 | GCTAGTGTCGTTGTCATACTTCATTTCTATTCG |
| LASO 33 | GCTTGCTTTATGTCTGTACAGAGATTAACGTTT |
| LASO 34 | CTGCTTGTCTTATGTATAAATGGAATGAATGAA |
| LASO 35 | CACTCTTACTTCTGCATAACCTCTACTTACTCT |
| LASO 36 | CTCGTTTATCCCTGCATATCTTATCTTATCTAT |
| LASO 37 | CGTTAGTTTATGTTGCCGATCAATTTCTATCTT |
| LASO 38 | CGTCTTTGTATGTCTGTAACAATAAGTGATTCG |
| LASO 39 | GCTTGTTCTGTTCCGATAGAAGGTAAGATTAAG |
| LASO 40 | GCTGATATGATGTAGTTACATACACTTTCGATG |
| LASO 41 | CGTTTCATTACCGTTAGAACCAATCTTTGTCCT |

| | |
|---|---|
| LASO 42 | TCGATTGTTATGTCTTGACATCAATTCATATTT |
| LASO 43 | CCATTGTCTTGCTCTATAATCTATTACTACTTA |
| LASO 44 | CCTGATTCTGACACGATACTTACTACTTTGACA |
| LASO 45 | AGGCTTGTTATAGTTATGACAATTCATTCATTT |
| LASO 46 | GCTTCTGTTAGTTTCATACAAATCAATTACATA |
| LASO targeting mRFP (variant 1) | GAACTCTTTGATAACGTCTTCGCTACTCGCATAGGTACCTT ATTCA |
| LASO targeting mRFP (variant 2) | GCATGAACTCTTTGATAACGTCTTCGCTACTCGCATAGGTA CCTTTTAATGAATTCA |
| LASO targeting mRFP (variant 3) | GCATGAACTCTTTGATAACGTCTTCGCTACTCGCATAGGTA CCTTAATGAATTCA |
| LASO targeting mRFP (variant 4) | GCATGAACTCTTTGATAACGTCTTCGCTACTCGCATAGGTA CCTTTGAATTCA |
| LASO targeting mRFP (variant 5) | GCATGAACTCTTTGATAACGTCTTCGCTACTCGCATAGGTA CCTTAATTCA |
| LASO targeting mRFP (variant 6) | GCATGAACTCTTTGATAACGTCTTCGCTACTCGCATAGGTA CCTTATTCA |
| LASO targeting mRFP (variant 7) | GCATGAACTCTTTGATAACGTCTTCGCTACTCGCATAGGTA CCTTTTCA |
| LASO targeting mRFP (variant 8) | GCATGAACTCTTTGATAACGTCTTCGCTACTCGCATAGGTA CCTTCA |
| LASO targeting mRFP (variant 9) | GCATGAACTCTTTGATAACGTCTTCGCTACTCGCATAGGTA CCTT |
| LASO targeting mRFP (variant 10) | GAACTCTTTGATAACGTCTTCGCTACTCGCATAGGTACCTTT TAATGAATTCA |
| LASO targeting mRFP (variant 11) | GAACTCTTTGATAACGTCTTCGCTACTCGCATAGGTACCTT AATGAATTCA |
| LASO targeting mRFP (variant 12) | GAACTCTTTGATAACGTCTTCGCTACTCGCATAGGTACCTTT GAATTCA |
| LASO targeting mRFP (variant 13) | GAACTCTTTGATAACGTCTTCGCTACTCGCATAGGTACCTT AATTCA |
| LASO targeting mRFP (variant 14) | GAACTCTTTGATAACGTCTTCGCTACTCGCATAGGTACCTTT TCA |
| LASO targeting mRFP (variant 15) | GAACTCTTTGATAACGTCTTCGCTACTCGCATAGGTACCTT CA |
| LASO targeting mRFP (variant 16) | GAACTCTTTGATAACGTCTTCGCTACTCGCATAGGTACCTT |
| LASO targeting mRFP (variant 17) | CTTTGATAACGTCTTCGCTACTCGCATAGGTACCTTTTAATG AATTCA |
| LASO targeting mRFP (variant 18) | CTTTGATAACGTCTTCGCTACTCGCATAGGTACCTTAATGAA TTCA |
| LASO targeting mRFP (variant 19) | CTTTGATAACGTCTTCGCTACTCGCATAGGTACCTTTGAATT CA |

| | |
|---|---|
| LASO targeting mRFP (variant 20) | CTTTGATAACGTCTTCGCTACTCGCATAGGTACCTTAATTCA |
| LASO targeting mRFP (variant 21) | CTTTGATAACGTCTTCGCTACTCGCATAGGTACCTTATTCA |
| LASO targeting mRFP (variant 22) | CTTTGATAACGTCTTCGCTACTCGCATAGGTACCTTTTCA |
| LASO targeting mRFP (variant 23) | CTTTGATAACGTCTTCGCTACTCGCATAGGTACCTTCA |
| LASO targeting mRFP (variant 24) | CTTTGATAACGTCTTCGCTACTCGCATAGGTACCTT |
| LASO targeting mRFP (variant 25) | ATAACGTCTTCGCTACTCGCATAGGTACCTTTTAATGAATTCA |
| LASO targeting mRFP (variant 26) | ATAACGTCTTCGCTACTCGCATAGGTACCTTTGAATTCA |
| LASO targeting mRFP (variant 27) | ATAACGTCTTCGCTACTCGCATAGGTACCTTAATTCA |
| LASO targeting mRFP (variant 28) | ATAACGTCTTCGCTACTCGCATAGGTACCTTATTCA |
| LASO targeting mRFP (variant 29) | ATAACGTCTTCGCTACTCGCATAGGTACCTTTTCA |
| LASO targeting mRFP (variant 30) | ATAACGTCTTCGCTACTCGCATAGGTACCTTCA |
| LASO targeting mRFP (variant 31) | GTCTTCGCTACTCGCATAGGTACCTTTTAATGAATTCA |
| LASO targeting mRFP (variant 32) | GTCTTCGCTACTCGCATAGGTACCTTAATGAATTCA |
| LASO targeting mRFP (variant 33) | GTCTTCGCTACTCGCATAGGTACCTTAATTCA |
| LASO targeting mRFP (variant 34) | GTCTTCGCTACTCGCATAGGTACCTTTTCA |
| LASO targeting mRFP (variant 35) | GTCTTCGCTACTCGCATAGGTACCTTCA |
| LASO targeting mRFP (variant 36) | GTCTTCGCTACTCGCATAGGTACCTT |
| LASO targeting mRFP (variant 37) | CGCTACTCGCATAGGTACCTTTTAATGAATTCA |
| LASO targeting mRFP (variant 38) | CGCTACTCGCATAGGTACCTTAATGAATTCA |
| LASO targeting mRFP (variant 39) | CGCTACTCGCATAGGTACCTTAATTCA |
| LASO targeting mRFP (variant 40) | CGCTACTCGCATAGGTACCTTATTCA |
| LASO targeting mRFP (variant 41) | CGCTACTCGCATAGGTACCTTTTCA |

| | |
|---|---|
| LASO targeting mRFP (variant 42) | CGCTACTCGCATAGGTACCTTCA |
| LASO targeting mRFP (variant 43) | CGCTACTCGCATAGGTACCTT |
| LASO targeting mRFP (variant 44) | CTCGCATAGGTACCTTAATGAATTCA |
| LASO targeting mRFP (variant 45) | CTCGCATAGGTACCTTTGAATTCA |
| LASO targeting mRFP (variant 46) | CTCGCATAGGTACCTTAATTCA |
| LASO targeting mRFP (variant 47) | CTCGCATAGGTACCTTATTCA |
| LASO targeting mRFP (variant 48) | CTCGCATAGGTACCTTTTCA |
| LASO targeting mRFP (variant 49) | CTCGCATAGGTACCTTCA |
| LASO targeting mRFP (variant 50) | CTCGCATAGGTACCTT |
| LASO targeting mRFP (variant 51) | CATGGTACCTTTCTCCTCTTTAATGAATTCA |
| LASO targeting mRFP (variant 52) | GCATGAACTCTTTGATAACGTCTTCGCTACTCGC |
| LASO targeting *gnd* | TGTTGTTTGGAATGTATATAGGTCGCCGTGTTGAA |
| LASO targeting *gmd* | GCGACTTTTGGCTAGTATTATGAATTTATTCAGTT |
| LASO targeting *murC* | TGTTGTGTATTATTTCTTTATGCCATTGAACGATG |
| LASO targeting *pfkA* | ATTTTCTTAATATGGACTAGATGCAAGGTGGAGTC |
| LASO targeting *zwf* | TGTGTTATCGTCTCGTCATCTAACCTGGTATTTAA |

**Supplementary Table 7.** Other target and trigger sequences used in this study.

| Description | Sequence |
|---|---|
| Genomic sfGFP 5' UTR (sfGFP CDS)[1] | TGAATTCATTAAAGAGGAGAAAGGTACCATGAGCAAAGGAGAAGAACTTTTCACTGGAGTTGTCCCAATTCTTGTTGAATTAGATGGTGATGTTAATGGG… |
| Genomic mRFP 5' UTR (mRFP CDS)[1] | TGAATTCATTAAAGAGGAGAAAGGTACCATGGCGAGTAGCGAAGACGTTATCAAAGAGTTCATGCGTTTCAAAGTTCGTATGGAAGGTTCCGTTAACGGT… |

| | |
|---|---|
| Antisense RNA (asRNA) targeting mRFP | ATAACGTCTTCGCTACTCGCCATGGTACCTTTCTCCTCTTTAATGAATTCA |
| Toehold switch 1 (including 21nt linker sequence) | AGAAAGAAGAAGAGATGCGGAAGAGAGAAATATAACACAAAGAGGAGAATATTTATGTCTTCCGCAAGACAACAAACAATAAACAGC |

**Supplementary Note 1.** Computational design of toehold repressors (design motif A) using NUPACK[2].

material = rna
temperature[C] = 37.0
trials = 10
sodium[M] = 1.0
dangles = some
allowmismatch = true

*#define structures (using DU+ or dot-bracket notation)*
structure TARGET = U15 D9 (U3 D6 (U15) U3) U51
structure TRIGGER = U33 U7 D10 (U4) U5
structure BOUND = D33 (U12 D9 (U3 D6 (U15) U3) U21 +) U7 D10 (U4) U5

*#sequence domains*
domain linear = N15
domain stem = N18
domain loop = N12
domain hairpin_2 = N25 agaggaga N6 atg N9
domain linker = aacctggcggcagcgcaaaag
#NOTE: designs 1, 3, 4, 6, 15, and 17 use a randomized linker (N21)
domain T500 = ggatctcaaagcccgccgaaaggcgggcttttttt

*#thread sequence domains*
TARGET.seq = linear stem loop hairpin_2 linker
TRIGGER.seq = stem* linear* T500
BOUND.seq = linear stem loop hairpin_2 linker stem* linear* T500

*#specify stop conditions*
TARGET.stop = 1
TRIGGER.stop = 1
BOUND.stop = 1

prevent = AAAA, GGGG, CCCC, UUUU, RRRRRR, YYYYYY, MMMMMM, KKKKKK, SSSSSS, WWWWWW

**Supplementary Note 2.** Computational design of toehold repressors (design motif B) using NUPACK[2].

material = rna
temperature[C] = 37.0
trials = 10
sodium[M] = 1.0
dangles = some
allowmismatch = true

*#define structures (using DU+ or dot-bracket notation)*
structure TARGET = U15 D12 (U2 D12 (U5) U2) U54
structure TRIGGER = U41 U7 D10 (U4) U5
structure BOUND = D41 (U10 D12 (U3 D6 (U12) U3) U21 +) U7 D10 (U4) U5

*#sequence domains*
domain linear = N15
domain stem = S3 N23
domain loop = N10
domain hairpin_2 = N18 S3 N4 agaggaga N6 atg N12
domain linker = aacctggcggcagcgcaaaag
domain T500 = ggatctcaaagcccgccgaaaggcgggcttttttt

*#thread sequence domains*
TARGET.seq = linear stem loop hairpin_2 linker
TRIGGER.seq = stem* linear* T500
BOUND.seq = linear stem loop hairpin_2 linker stem* linear* T500

*#specify stop conditions*
TARGET.stop = 1
TRIGGER.stop = 1
BOUND.stop = 1

prevent = AAAA, GGGG, CCCC, UUUU, RRRRRR, YYYYYY, MMMMMM, KKKKKK, SSSSSS, WWWWWW

**Supplementary Note 3.** Design of a toehold switch for simultaneous activation and repression[2,3].

*#NOTE: This script designs a toehold switch to respond to trigger #1 for simultaneous activation and repression (Figure 7). Alternatively, the toehold switch and toehold repressor targets could be designed from a specified trigger sequence, or all three sequences (trigger, repressor target, and toehold switch target) could be designed together.*

material = rna
temperature[C] = 37.0
trials = 10
sodium[M] = 1.0
dangles = some
allowmismatch = true

*#define structures (using DU+ or dot-bracket notation)*
structure TARGET = U15 D9 (U3 D6 (U15) U3) U21
structure BOUND = D33 (U7 U8 U18 U21 +) U7 D10 (U4) U5

*#sequence domains*
domain linear = agaaagaagaagaga
domain 5prime = N18
domain loop = aacacaaagaggaga
domain 3prime = N6 atg N9
domain linker = agacaacaaacaataaacagc
domain trigger = atattttttcttccgcatctcttcttctttctggatctcaaagcccgccgaaaggcgggcttttttt

*#thread sequence domains*
TARGET.seq = linear 5prime loop 3prime linker
BOUND.seq = linear 5prime loop 3prime linker trigger

*#specify stop conditions*
TARGET.stop = 1
BOUND.stop = 1

prevent = AAAA, GGGG, CCCC, UUUU, RRRRRR, YYYYYY, MMMMMM, KKKKKK, SSSSSS, WWWWWW

### SUPPLEMENTAL FIGURES

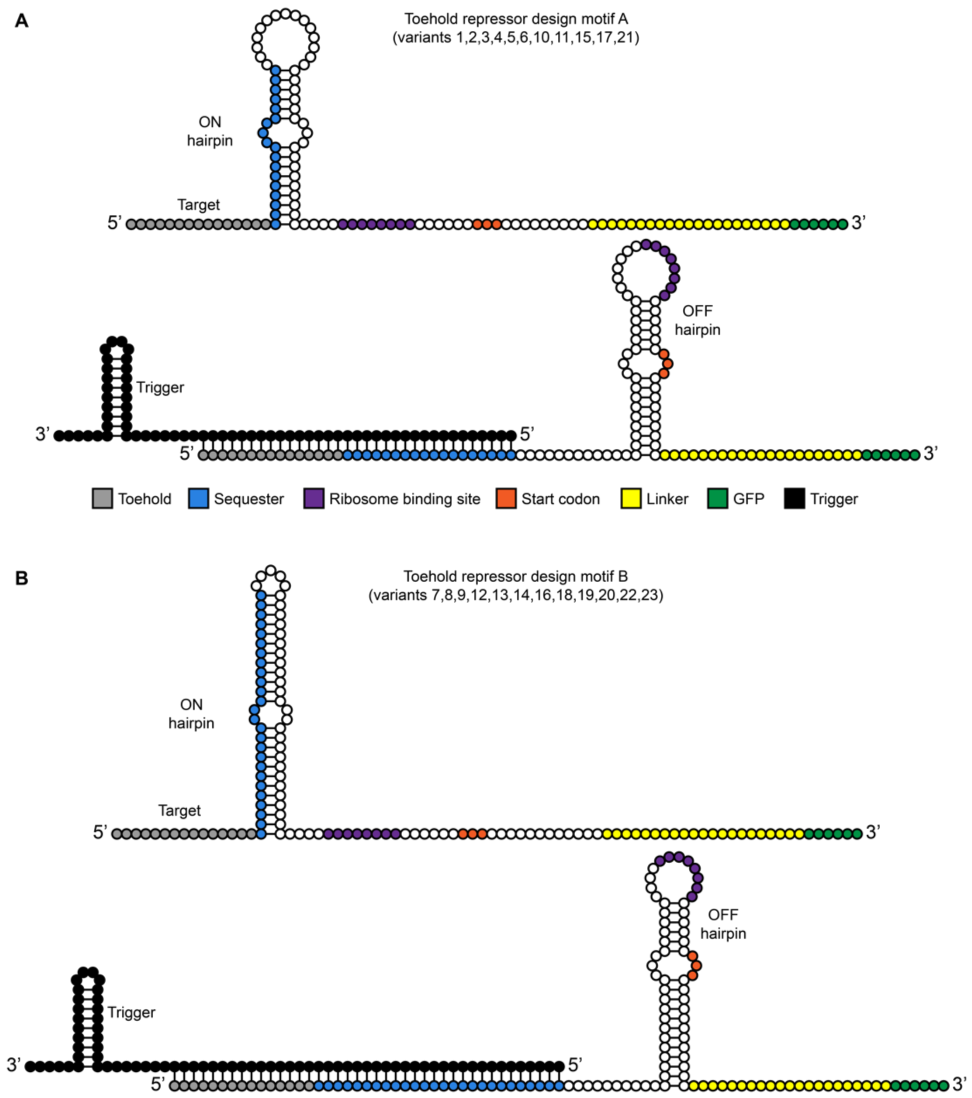


**Figure S1.** Toehold repressor design motifs used in this study. (A) Design motif A uses ON and OFF hairpin structures with identical secondary structures, which are based on the structure of the best-performing toehold switch translational activator designs from Green et al.[3]. (B) Design motif B uses a more stable OFF hairpin, with the goal of minimizing leak in the OFF state (as demonstrated by Pardee et al.[4]). To compensate for this more stable OFF hairpin, and to help drive formation to the ON state in the absence of a trigger RNA, a longer ON hairpin with a small loop was used. Each circle corresponds to a single nucleotide with colour coding indicated.

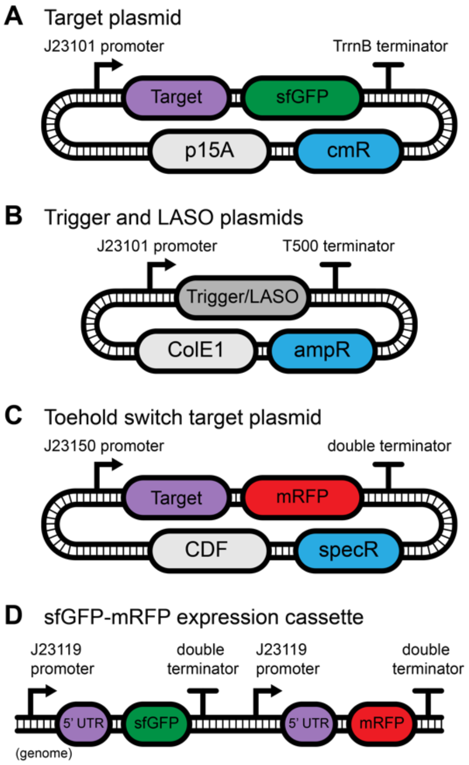


**Figure S2.** Schematics of plasmids used in this study. (A) Target RNAs are expressed from a moderately strong $\sigma^{70}$ promoter (J23101) and are followed by sfGFP and a double terminator from the *rrnB* operon (TrrnB). Target plasmids harbour a chloramphenicol resistance gene, cmR, and p15A, a medium copy number origin of replication. (B) Trigger RNAs and LASOs also use the J23101 promoter, but use a compact T500 terminator, a high copy number origin of replication (ColE1), and an ampicillin resistance gene, ampR. (C) To co-express a toehold repressor and toehold switch (Figure 7), the toehold switch target sequence was cloned upstream of mRFP in a plasmid harbouring a spectinomycin resistance gene, specR, and a CDF origin of replication. The copy number of the CDF plasmid is higher than p15A, so the strength of the toehold switch promoter was weakened to help balance expression of the two RNAs. (D) sfGFP and RFP expression cassette (Figure 4). Both sfGFP and mRFP are expressed by strong J23119 promoters at a genomic locus using the same 5' UTR sequence. Promoters were selected based on a previous characterization study[5], and all sequences were obtained from the iGEM Registry of Standard Biological Parts (http://parts.igem.org).

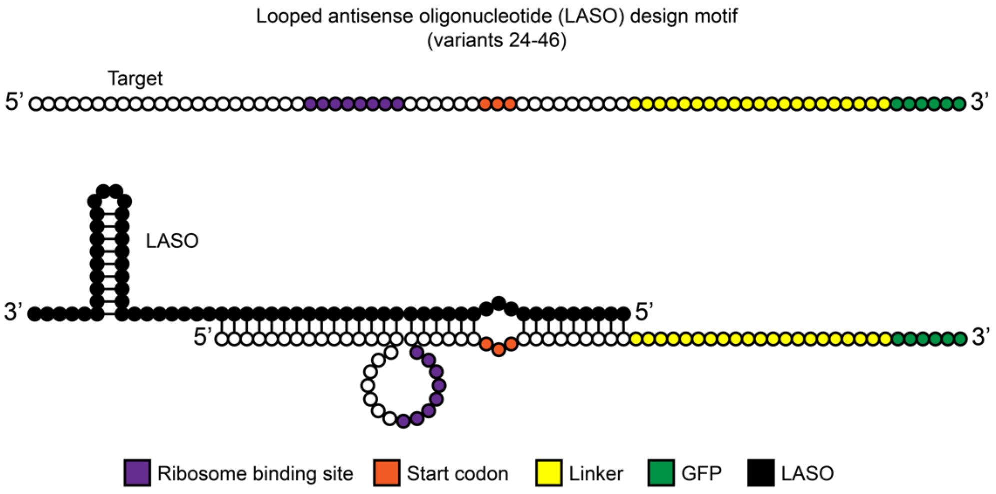


**Figure S3.** The LASO design motif used in this study. (Top) The target mRNA colour coded with relevant sequence domains. (Bottom) The designed interaction between the LASO RNA and the target showing how the RBS and AUG sequence domains become 'looped' upon binding. Each circle corresponds to a single nucleotide with colour coding indicated.

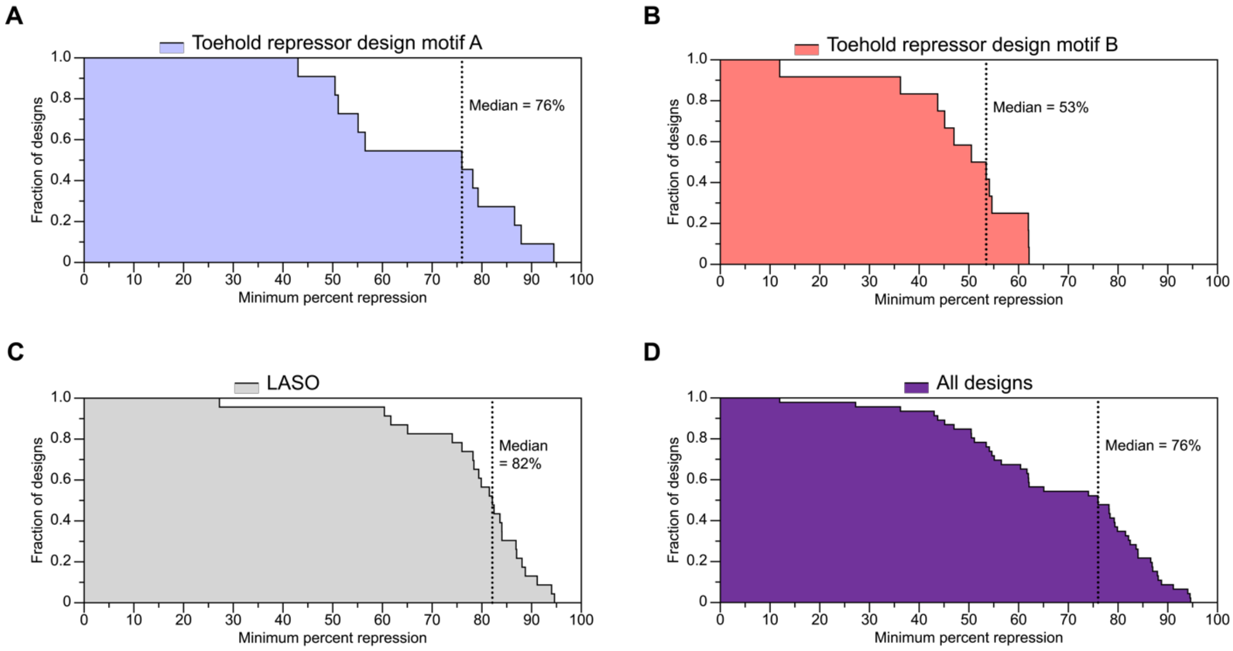


**Figure S4.** Comparing the performance of the design motifs pursued in this study. Each plot shows the fraction of designs tested (Y-axis) which repress translation at or above a given threshold (X-axis). Median repression for each set of designs is also indicated. (A) Toehold repressor design motif A showed better repression compared to design strategy B (B), with 76% and 53% median repression, respectively. (C) The LASO designs showed the best performance, with median repression of 82%. (D) Median repression of 76% was observed for all designs.

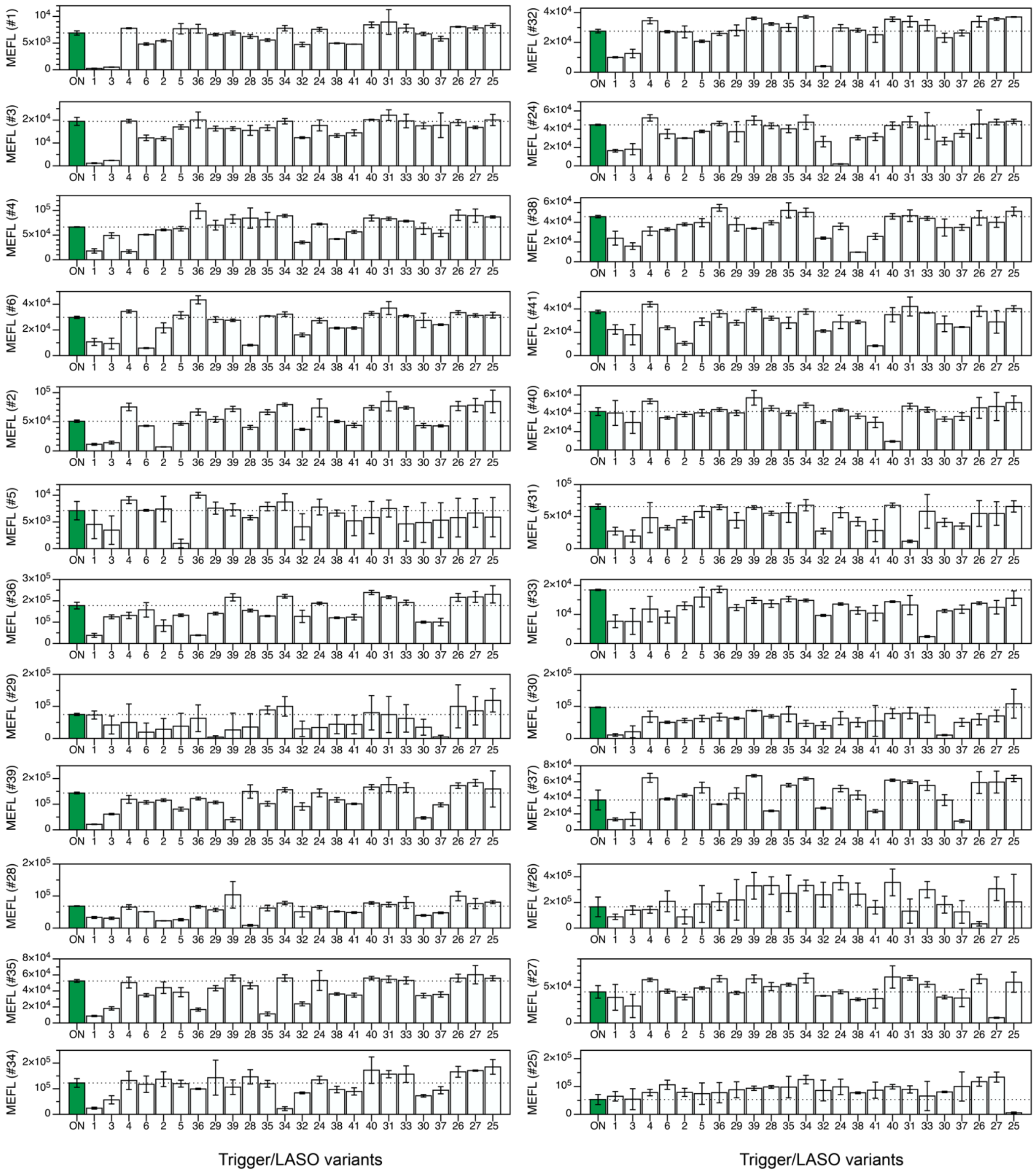


**Figure S5.** Expression data used to construct the orthogonality matrix shown in Figure 2. Each plot corresponds to a single column in Figure 2. No trigger (ON) expression from the target is shown in green, and expression with each trigger or LASO variant is shown in white. Error bars correspond to the standard deviation of at least three biological replicates.

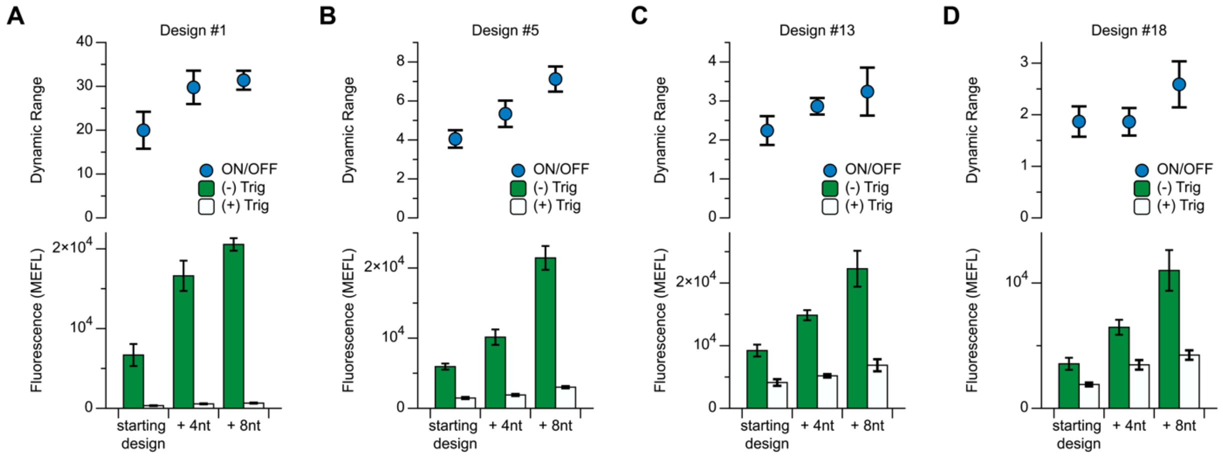


**Figure S6.** Reducing overlap between ON hairpin and ribosome footprint improves both ON level and dynamic range in several toehold repressor designs. Design #1 (A), design #5 (B), design #13 (C), and design #18 (D) (from Figure 1C) have increased ON fluorescence when 4nt or 8nt are added before the RBS. In all cases, this increase in ON fluorescence corresponds to an increase in dynamic range.

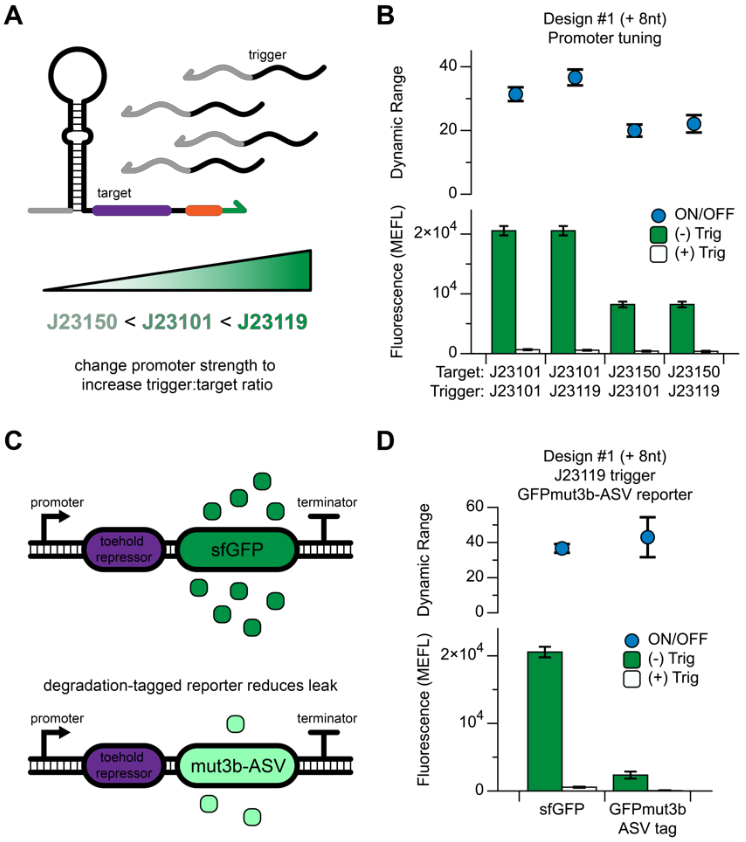


**Figure S7.** The effect of promoter tuning and reporter protein stability on the dynamic range of toehold repressor design #1 (from Figure 1C). (A) The ratio of trigger to target molecules can be increased by increasing the strength of the trigger promoter and/or decreasing the strength of the target promoter. (B) Promoter combinations were tested using design #1 (including 8nt added before the RBS). The best-performing combination was the moderate strength promoter (J23101) on the target and a strong promoter (J23119) on the trigger. This design was then tested with GFPmut3b-ASV, a degradation-tagged GFP variant (C). This reduced both ON and OFF levels with insignificant changes to the dynamic range (D).

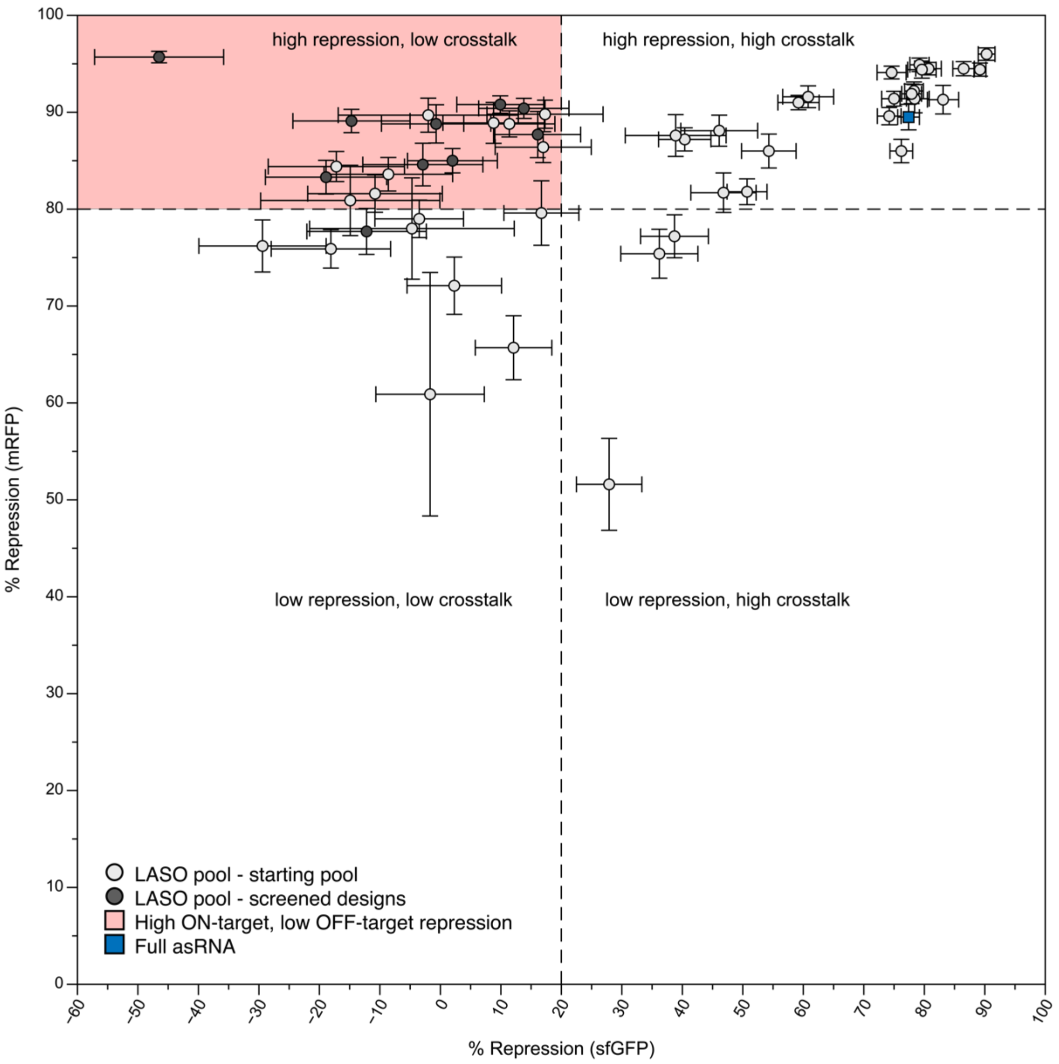


**Figure S8.** Repression plot of the library of LASOs targeting mRFP and sfGFP shown in Figure 4, including uncertainty in measured repression values. Repression of mRFP (Y-axis) and sfGFP (X-axis) is plotted for each LASO design (grey circles) as well as the full asRNA (blue square).

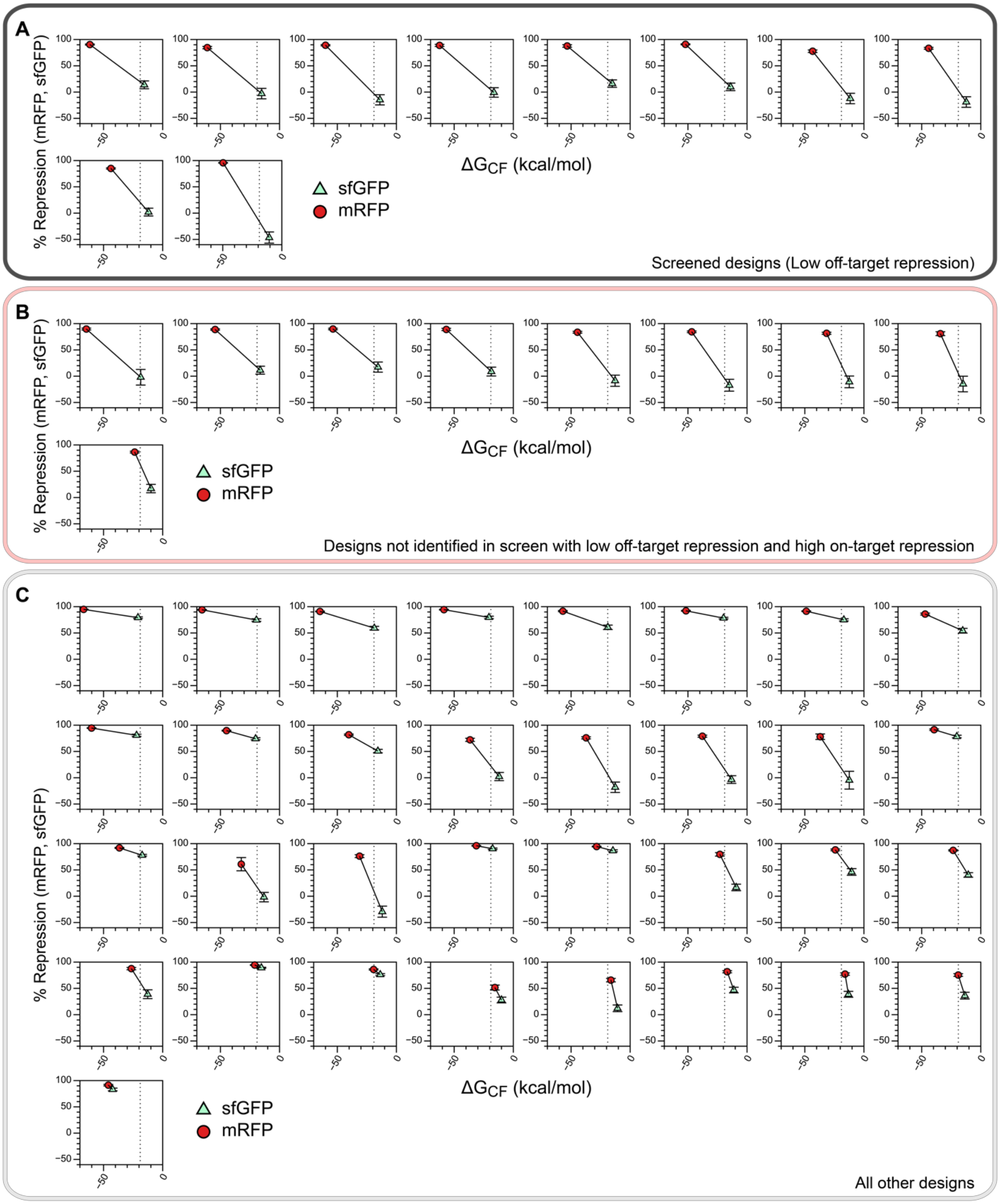


**Figure S9.** Repression vs. $\Delta G_{CF}$ for all LASO designs targeting mRFP from Figure 4F. (A) Designs identified in screen (Figure 4H) to have low off-target repression of sfGFP. (B) Designs with low measured off-target repression (sfGFP) and high on-target repression (mRFP) which were not identified from the screening workflow. (C) All other designs, which display low on-target and/or high off-target repression. Vertical dashed lines indicate a $\Delta G_{CF}$ value of -19 kcal/mol.

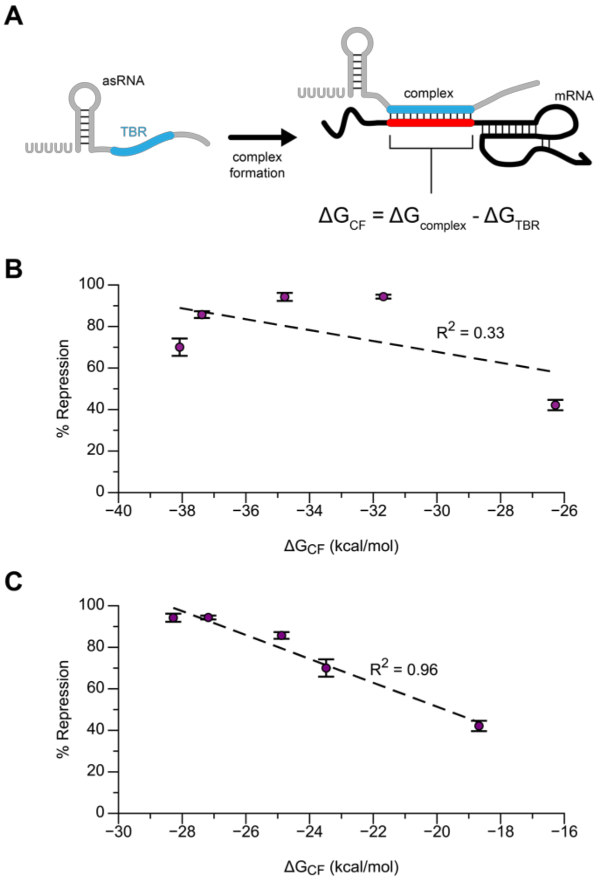


**Figure S10.** Updated definition of $\Delta G_{CF}$ displays improved correlation with LASO repression efficiency of endogenous mRNA sequences. (A) Definition of $\Delta G_{CF}$ proposed by Hoynes-O'Connor and Moon[6], which compares the free energy of the asRNA-mRNA duplex subregion ($\Delta G_{complex}$) to the free energy of the mRNA-binding subsequence of the asRNA alone (termed the target binding region, $\Delta G_{TBR}$). (B) This original definition shows a moderate correlation with repression efficiency of LASO designs shown in Figure 5 ($R^2$ = 0.33). (C) The updated definition of $\Delta G_{CF}$ proposed in this work better captures differences in LASO repression efficiency ($R^2$ = 0.96), likely because this metric accounts for the effects of intramolecular structures formed by the mRNA target.

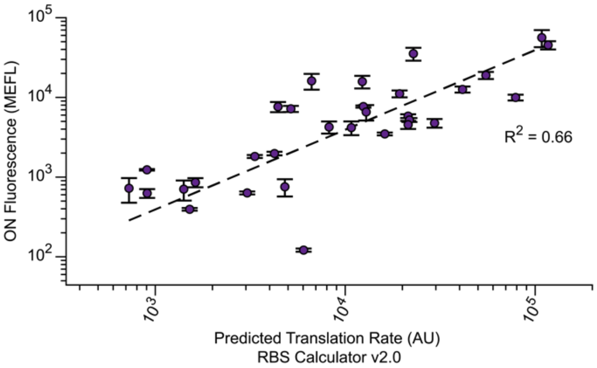


**Figure S11.** Expression levels for library members from a randomized RBS screen (Figure 6) correlate with predicted translation rate calculated using RBS Calculator v2.0.

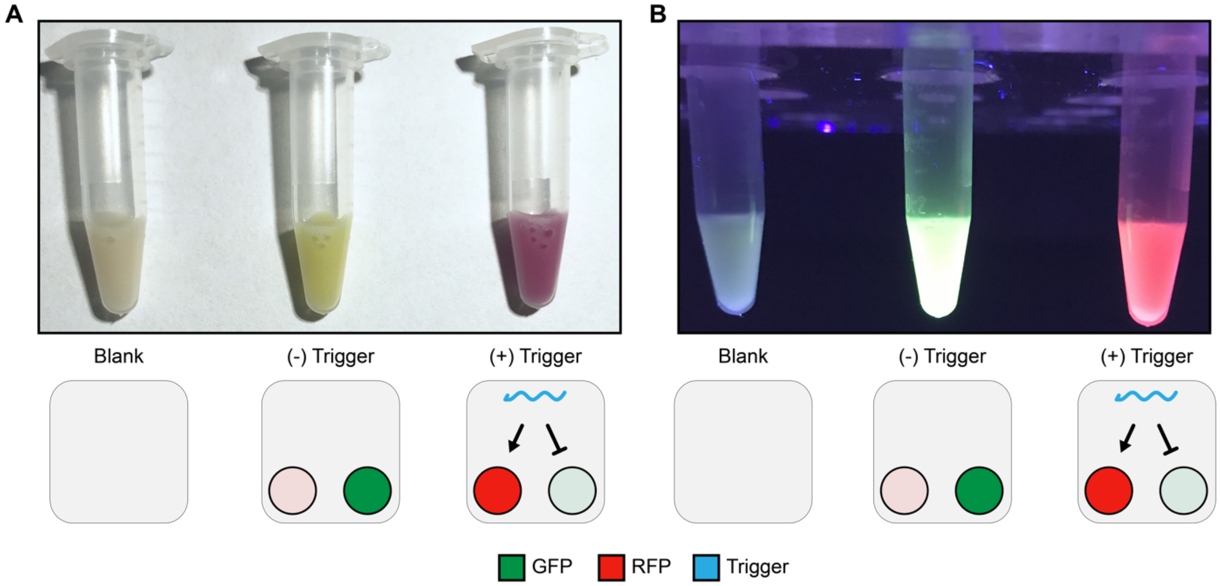


**Figure S12.** Simultaneous activation and repression from a toehold repressor and toehold switch (Figure 7) can be visualized by eye. 5 mL of cells containing blank control plasmids (blank), target plasmids (- trigger), or target plasmids with trigger (+ trigger) were grown overnight in LB then resuspended in 100 µL PBS. Switching of expression from sfGFP (- trigger) to mRFP (+ trigger) is clearly visible by eye (A) and under UV illumination (B).
